# Diversity in Rubisco Kinetics and CO₂-Concentrating Mechanisms Among Cyanobacterial Lineages

**DOI:** 10.64898/2025.12.30.696979

**Authors:** Pere Aguiló-Nicolau, Reto S Wijker, Concepción Iñiguez, Sebastià Capó-Bauçà, Heather M Stoll, Jeroni Galmés

## Abstract

Cyanobacteria are the most ancient oxygenic photosynthetic organisms on Earth and play a pivotal role in the global carbon cycle. Despite their ecological and evolutionary significance, the mechanisms of carbon acquisition and fixation in this phylum remain largely unexplored beyond a few model species. Here, we examined representative taxa spanning the full phylogenetic breadth of Cyanobacteria, assessing *in vivo* carbon-acquisition pathways, the role and effectiveness of CO_2_-concentrating mechanisms (CCMs), as well as conducting *in vitro* biochemical characterizations of the kinetic traits and carbon isotope fractionation of Rubisco. We found significant lineage-specific differences in Rubisco kinetics and CCM performance, but a common signature of high Rubisco catalytic turnover coupled with low CO_2_ affinity—consistent with the co-evolution of this enzyme together with powerful CCMs. Furthermore, we identified a strong positive correlation between Rubisco carbon isotope fractionation and its CO_2_/O_2_ specificity factor. Together, these results provide fresh insight into Rubisco catalysis and shed light on its co-evolution with CCMs, underscoring their role in shaping Earth’s carbon dynamics.

## Introduction

Cyanobacteria are among the most ecological and evolutionary relevant organisms on Earth. They contribute ∼25% of global inorganic carbon fixation and support nearly half of marine primary production (Mishra, 2020). Their significance extends deep into Earth’s history: ∼3.5 billion years ago (Ga), Cyanobacteria evolved oxygenic photosynthesis, releasing molecular oxygen and triggering the Great Oxidation event (GOE), which dramatically reshaped Earth’s atmosphere and enabled the emergence of eukaryotic life (Sánchez-Baracaldo et al., 2022).

Central to oxygenic photosynthesis is ribulose-1,5-bisphosphate carboxylase oxygenase (Rubisco), the primary CO_2_-fixing enzyme of the Calvin–Benson–Bassham (CBB) cycle, which underpins nearly all trophic networks. Rubisco likely originated from a non-CO_2_-fixing ancestral enzyme in a CO_2_-rich anoxygenic environment ∼3.5 Ga and diversified into the present-day forms (I, II, III, II/III) through horizontal gene transfers and structural adaptations (Tabita et al., 2008). However, the rise of atmospheric O_2_ exposed a fundamental limitation for its catalysis: Rubisco catalyzes both the carboxylation and the oxygenation of Ribulose biphosphate, with the latter triggering photorespiration and causing net loss of fixed carbon (Bauwe et al., 2012).

Rubisco inefficiency to fully discriminate between CO_2_ and O_2_, combined with low CO_2_ availability in aquatic systems, drove the evolution of CO_2_-concentrating mechanisms (CCMs) in Cyanobacteria and other autotrophs between ∼1.8 and ∼0.7 Ga (Hagemann et al., 2021). Cyanobacterial biophysical CCMs include bicarbonate transporters, thylakoid CO_2_-uptake complexes, ion-gradient systems, and specialized microcompartments—carboxysomes—that encapsulate Rubisco and carbonic anhydrases (CAs), raising the local CO_2_/O_2_ ratio around Rubisco active sites (Hagemann et al., 2021). Two types of Cyanobacteria can be classified according to their CCM arrangement: β-cyanobacteria present β-carboxysomes, with form IB Rubisco and a higher diversity of bicarbonate transporters and CO_2_-uptake systems compared to α-cyanobacteria, which present α-carboxysomes with form IA Rubisco (Long et al., 2016). CCMs not only underpin cyanobacterial photosynthesis but are also inspiring synthetic biology approaches to improve crop productivity (Nguyen et al., 2024).

Cyanobacterial Rubisco kinetics reflect the interplay between ancient biochemical constraints and CCMs. Compared with C_3_ land plants, cyanobacterial Rubisco generally exhibits lower CO_2_ specificity (S_c/o_) but higher carboxylation turnover rates (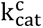), consistent with operation in a high-CO_2_ microenvironment (Savir et al., 2010). For instance, *Synechococcus* sp. PCC 6301 Rubisco exhibits S_c/o_ ≈ 44 mol mol^-1^ and 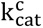 ≈ 10 s^-1^, while C_3_ plant Rubisco is slower 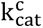 ≈ 3 s^-1^ but more CO_2_-specific (S_c/o_ ≈ 100 mol mol^-1^), reflecting adaptation to low CO_2_ and high O_2_ in absence of CCMs (Iñiguez et al., 2020). Across Cyanobacteria, reported Rubisco kinetic traits vary widely—S_c/o_ 32-60 mol mol^-1^, half-saturation constant for CO_2_ (K_c_) 80-309 μM, half-saturation constant for O_2_ (K_o_) 529-1,400 μM, and 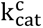 2.4-14.4 s^-1^—yet these values are based largely on a handful of model strains (de Pins et al., 2025). Only recently non-model species have been examined, with polyextremophiles revealing exceptional Rubisco performance (Aguiló-Nicolau et al., 2023). Such findings suggest that the true enzymatic diversity within Cyanobacteria remains underexplored.

Rubisco also describes a kinetic isotope effect (KIE), preferentially fixing ^12^CO_2_ over ^13^CO_2_, thereby producing ^13^C-depleted biomass. This intrinsic isotope fractionation (ε_Rubisco_) has been proposed to correlate with S_c/o_, as both parameters are governed by transition state chemistry (Tcherkez et al., 2006). The preferential fixation of ^12^CO_2_ arises from its higher vibrational frequency, which lowers the activation energy for bond formation. In addition, a closer resemblance between the carboxylation transition state and its product further amplifies fractionation (Tcherkez et al., 2006). A similar mechanism underlies Rubisco’s discrimination between CO_2_ and O_2_ (Tcherkez et al., 2006). However, the potential trade-of between ε_Rubisco_ and S_c/o_ remains unresolved.

Exploring the natural diversity of ε_Rubisco_ in broader phylogenetic groups, such as Cyanobacteria, is particularly relevant because ε_Rubisco_ values are used in reconstructions of past atmospheric CO_2_ levels and global carbon cycling (Wang et al., 2023b). Although most models assume a constant ε_Rubisco_ of ∼29‰, empirical values vary widely—from ∼11‰ in coccolithophores to ∼30‰ in higher plants (Wang et al., 2023a). Constraining ε_Rubisco_ is therefore essential for interpreting Earth’s geological history, particularly in the Precambrian, when Cyanobacteria were the predominant primary producers recorded in the fossil record (Garcia et al., 2021).

Despite Cyanobacteria ecological and evolutionary significance, our understanding of their carbon acquisition mechanisms across the different lineages of the group remains incomplete. Current Rubisco kinetic datasets underrepresent the enzymatic diversity of the phylum, while both the variability of ε_Rubisco_ and its relationship with S_c/o_ remain unresolved. In addition, the diversity of CCMs and their co-evolution with Rubisco are still poorly understood.

Here we address these gaps by testing whether phylogenetically and ecologically diverse Cyanobacteria exhibit distinct photosynthetic traits, CCM efficiencies, and Rubisco kinetics that reflect both evolutionary history and environmental adaptation. We hypothesize that species from contrasting lineages and niches have evolved different carbon acquisition pathways, detectable across physiological, biochemical, and ultrastructural levels. To this end, we performed a comparative characterization of five cyanobacterial strains representative of the main lineages, integrating anatomical analyses, photosynthetic performance, CCM function, Rubisco kinetics, and ε_Rubisco_ determinations. This comprehensive approach provides new insights into the diversification of cyanobacterial carbon assimilation with implications for global biogeochemical models, paleoenvironmental reconstructions, and synthetic biology.

## Materials and methods

### Cyanobacterial Culturing, Biomass Production and Harvesting

Strains were selected based on the phylogenetic analysis in Komárek et al. (2014) and the availability of fully sequenced genomes in GenBank, except for *Chroococcidiopsis thermalis*, included for its extremotolerant capacity. Cultures were obtained from the Pasteur Culture Collection of Cyanobacteria (PCC, France) and the Culture Collection of Autotrophic Organisms (CCALA, Czech Republic). The selected species included a representative strain of each major β-cyanobacterial lineage: *Oscillatoria accuminata* (PCC 6304), representing the order Oscillatoriales; *Cyanobacterium aponinum* (PCC 10605), from Chroococcales; *Gloeobacter violaceus* (PCC 7421), representing Gloeobacterales; *Synechococcus* sp. (PCC 6301), from Synechococcales, and *C. thermalis* (KOMAREK 1964/111), from Chroococcidiopsidales.

All the strains were grown in 50 or 250 mL sterile, ventilated cell culture flasks (CCFP-25V-100, Labbox, Spain). *Cyanobacterium aponinum* and *Synechococcus* sp. were grown under continuous agitation, whereas *G. violaceus*, *O. acuminata,* and *C. thermalis* were maintained under static conditions. BG11 medium (Waterbury and Stanier, 1981) was used to grow *G. violaceus*, *O. acuminata* and *C. aponinum*, while *S. elongatus* and *C. thermalis* were grown in Z-medium (Andersen, 2005). All cultures were maintained at 25 °C under a 16:8 light-dark cycle (Aralab Fitoclima S600, PLH, Spain), and a light intensity of 50 μmol m^-2^ s^-1^ provided by 4,000 K fluorescent tubes (Osram L 18W/840 Lumilux, Germany). Growth was monitored spectrophotometrically at 650 nm (OD_650_; Multiskan Sky 1530-00433C, Thermo Scientific, USA). Once OD_650_ reached approximately 0.5, biomass was harvested, and cultures were diluted with fresh medium.

### *In vivo* Measurements

Clark-type oxygen electrode chambers (Oxygraph+ system, Hansatech, UK) were used to conduct all the *in vivo* measurements at 25 °C, including photosynthesis-C curves and net photosynthetic rates with and without the effects of inhibitors targeting internal and external carbonic anhydrases (CAs) as well as anion-exchange bicarbonate transporters.

Photosynthesis-C curves were performed by gently centrifuging 4 mL of cultures (OD_650_ ≈ 0.5) at 3,000 x *g* for 3 min at 25 °C, followed by washing of the pellet and resuspension in 2 mL of CO_2_-free 20 mM Tris-buffered BG11 or Z-medium, depending on the strain. The addition of Tris buffer had no detectable effect on the photosynthetic rate in any of the cyanobacterial strains (data not shown). The resuspended culture was transferred to the oxygen electrode chamber under saturating white light (300 μmol m^-2^ s^-1^) and continuous magnetic stirring. Measurements began in CO_2_-free medium, and O_2_ evolution was monitored using OxiTrace+ software (version 1.0.48, Hansatech Instruments, UK). Steady-state conditions—defined by the absence of net O_2_ production—were typically reached within the first 30 min. Increasing concentrations of dissolved inorganic carbon (DIC), ranging from 0 to 1,500 μM, were sequentially added to the chamber. No significant pH changes were detected during the measurements. O_2_ evolution rates were recorded every 3 min following stabilization after each DIC addition. Net O_2_ production rates were normalized to dry biomass weight (DW), by freeze-drying the 2 mL assay volume. Oxygen saturation levels in air-equilibrated media at 25 °C were determined using DOTABLES (http://water.usgs.gov/software/DOTABLES/) at the corresponding medium conductivity. The equilibrium concentration of dissolved CO_2_—[CO_2_]_aq_—corresponding to each DIC addition was calculated using CO2SYS software (v.2.2). The maximum photosynthetic rate (A_n max_) and the *in vivo* semi-saturation constant for CO_2_ (K_m *in vivo*_) were obtained by fitting the photosynthesis-C curves to the Michaelis–Menten equation, as described in Aguiló-Nicolau et al. (2023).

The inhibitory effects on the net photosynthetic rates caused by CCM inhibitors were assessed using 2 mL of culture (OD_650_ ≈ 0.5) placed in the oxygen electrode chamber under saturating white light (300 μmol m^-2^ s^-1^) and continuous stirring. The extracellular CA inhibitor acetazolamide (AZ) was added to a final concentration of 200 μM, and the change in O_2_ evolution rate was measured after 3 minutes of stabilization. Subsequently, both external and internal CAs inhibitor ethoxyzolamide (EZ) was added to the same chamber at a final concentration of 200 μM, and the response was again recorded. The effect of the anion-exchange inhibitor 4,4′-diisothiocyanatostilbene-2,2′-disulfonate (DIDS; 300 μM final concentration) was equally tested using a separate sample.

## Anatomical Characterizations

Transmission electron microscope (TEM) images were obtained by centrifuging 1 mL of culture (OD_650_ ≈ 0.5) at 8,000 x *g* for 3 min at 25 °C. The pellet was resuspended in 1 mL fixation buffer consisting of 0.1 M phosphate buffer (pH 7.2) containing 4% glutaraldehyde and 2% paraformaldehyde. Post-fixation was performed in 1% osmium tetroxide, prepared in 0.1 M Sorensen’s phosphate buffer, for 1 h as described in Aguiló-Nicolau et al. (2023). Fixed samples were stained in 2% uranyl acetate, dehydrated in a graded ethanol series, and embedded in London Resin White (EMS, USA).

Semithin (1 µm thick) and ultrathin (50 to 70 nm thick) sections were obtained using an ultramicrotome (UC7/FC7, Leica, Germany). Semithin sections were mounted on glass slides and stained with epoxy tissue stain (EMS, USA). The ultrathin sections were mounted on copper grids and visualized using a TEM (JEM 1400, Jeol Ltd., Japan). Image analysis was performed using ImageJ software (Wayne Rasband National Institutes of Health, version 1.54p).

## Extraction and Quantification of Total Soluble Proteins and Rubisco

A volume of 50 to 150 mL of culture (OD_650_ ≈ 0.5) was harvested by centrifugation at 8,000 x *g* for 3 min at 25 °C, snap-frozen and stored at -80 °C. The pelleted cells where resuspended in 4 mL cold extraction buffer containing 100 mM bicine (pH 8.1), 20 mM MgCl_2_, 1 mM EDTA, 1 mM benzamidine, 1 mM ε-aminocaproic acid, 10 mM dithiothreitol (DTT), 2% plant protease inhibitor cocktail (P9599, Merck, USA), 100 mM β-mercaptoethanol, 20 mM phenylmethanesulfonylfluoride (PMSF), 2% CelLytic^TM^ B (B7435, Merck, USA), 2.5 mL of 2 mm glass beads, and 0.1 g of polyvinylpolypyrrolidone (PVPP). Cell lysis was performed on an ice bath using a probe sonicator (UP200St, Hielscher Ultrasonics, Germany) for 5 min with 30 s (on/of) intervals at 40% amplitude. The homogenate was centrifuged at 8,000 x *g* for 15 min at 4 °C. The supernatant was aliquoted and stored at -80 °C.

An aliquot of the supernatant was used to quantify the total soluble protein (TSP) content by Bradford’s method (Bradford, 1976). Rubisco active site concentration was quantified by incubating an aliquot of the supernatant with 25 mM NaHCO_3_ for 30 min at 25 °C for optimal Rubisco activation. This was followed by a 30 min incubation with 0.2 mM of ^14^C-labelled 2’-carboxy-D-arabinitol-1,5-bisphosphate (^14^C-CABP)—a specific Rubisco binding inhibitor (Ruuska et al., 1998). The optimal concentration of ^14^C-CABP for Rubisco quantification was determined individually for each strain as described in Capó-Bauçà et al. (2022)—data not shown. Unbound ^14^C-CABP was separated from Rubisco-bound ^14^C-CABP via Sephadex G-50 Fine column chromatography (17-0042-01, GE Healthcare, USA). The radioactivity of the fraction containing Rubisco-bound ^14^C-CABP was measured using a liquid scintillation counter (Tri-Carb 4910 TR, Revity, USA).

## *In vitro* Rubisco Kinetic Parameters Determination

An aliquot of the crude protein extract was partially purified using two 5 mL anion-exchange cartridges (Bio-Scale Mini Macro-Prep High Q Cartridge 7324124, Bio-Rad, USA) connected in series to a fast protein liquid chromatography system (ÄKTA pure^TM^ 25, Cytiva, USA). The eluate was subsequently desalted and concentrated approximately tenfold by centrifugation at 2,000 × *g* and 4 °C using a 10 kDa molecular weight cut-of filter (Amicon Ultra 4, UFC8010, Merck, USA).

Rubisco carboxylation kinetic parameters were measured at 25 °C. Assays were performed in 7 mL septum-capped crystal vials under magnetic stirring, each containing 0.4 mL of assay buffer (100 mM bicine pH 8.1, 20mM MgCl_2_, and 100 W-A units of CA, C3934, Merck, USA). The buffer was pre-equilibrated for 2 h with either 100% N_2_ or CO_2_-free synthetic air (21% O_2_, 79% N_2_). Eight concentrations of NaH^14^CO_3_ (0 to 60 mM; specific activity 3.7 x 10^10^ Bq · mol^-1^) and 1.6 mM RuBP—synthesized and purified as in Kane et al. (1994)—were added to each vial. Semi-purified extracts were preactivated by incubation with 20 mM NaH^14^CO_3_ at 35 °C for 20 min. The reaction was initiated by adding 20 µL of preactivated extract to the septum-capped crystal vials and allowed to proceed for 2 min before quenching with 200 µL of 10 M formic acid. Acid-stable, non-volatile ^14^C-labeled organic products were quantified by drying the reaction at 80 °C for 2 h, resuspending the residue in MilliQ water, and measuring radioactivity with a scintillation counter (Tri-Carb 4910 TR, Revity, USA).

The Michaelis–Menten equation was used to fit the data and estimate the CO_2_ semi-saturation constants at 0% and 21% O_2_ (K_c_ and 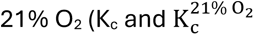, respectively), as well as the maximum carboxylation rate 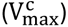. The semi-saturation O_2_ constant (K_o_) was determined by linear fitting between K_c_ and 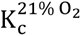. [CO_2_]_aq_ was calculated assuming a carbonic acid dissociation constant (pK_a_) of 6.11, calculated following Galmés et al. (2016), a solubility constant for CO_2_ of 0.034 mol L^-1^ atm^-1^ at 25 °C and accurate pH measurements of the assay buffer. The carboxylation turnover rate (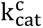) was determined by dividing 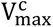by Rubisco active sites concentration, quantified using ^14^C-CABP binding assay, assuming eight active binding sites per Rubisco.

Rubisco specificity factor (S_c/o_) was determined in the semi-purified extracts following the method of Kane et al. (1994). A volume of 20-40 µL of extract was added to 7 mL septum-capped crystal vials containing 940 µL of assay buffer (30 mM triethanolamine-acetate, pH 8.3; 15 mM Mg-acetate), and 400 W-A units of CA. The buffer was equilibrated for 1 h with a gas mixture of 99.95% O_2_ and 0.05% CO_2_. The reaction was initiated by adding 1 nmol [1-^3^H] RuBP and allowed to proceed for 1 h at 25 °C under magnetic stirring and a continuous gas flow. The reaction was terminated with 0.35 U of alkaline phosphatase (P7640, Merck, USA) and purified using AG1-X8 anion-exchange resin (1401441, Bio-Rad, USA). The unphosphorylated products of carboxylation and oxygenation (glycerate and glycolate, respectively) were separated by HPLC (Jasco-UV-4075, Jasco inc., USA) and quantified by scintillation counting (Tri-Carb 4910 TR, Revity, USA). The S_c/o_ values were calculated using a CO_2_/O_2_ solubility ratio of 0.038 at 25 °C.

Rubisco kinetic parameters from *Synechococcus* sp. (PCC6301) and *Chroococcidiopsis thermalis* (KOMAREK 1964/111) were obtained from Aguiló-Nicolau et al. (2023). Other cyanobacterial lineages Rubisco kinetic parameters were extracted from the compilation by Iñiguez et al. (2020).

## *In vitro* Rubisco Carbon Isotope Fractionation

Rubisco carbon isotope discrimination (ε_Rubisco_) was determined using a modified method based on Scott et al. (2004), with particular refinements specified in Wijker et al. (2025) which provides results for *Synecococcus* in the range of previously published determinations but improved transparency in the precision of the determination.

In summary, for each assay, 88.5 mM EPPS (pH 7.8), 28.7 mM MgCl_2_, 0.5 mM NaHCO_3_, and 400 W-A units of CA were loaded into 10 mL gas-tight syringes (81656 Hamilton, USA), along with semi-purified Rubisco extract preactivated with 20 mM NaHCO_3_ at 35 °C for 15 min. After another 15 min of incubation at 25 °C, an initial aliquot (0.5 mL) was collected, diluted 1:5 with 110 mM EPPS (pH 7.8) and filtered using 10 kDa centrifugal filter (Amicon Ultra 4, UFC8010, Merck, USA) at 2,000 x *g* for 3 min at 25 °C. The resulting flow through was analyzed with a DIC-δ^13^C analyzer (AS-D1 and G2131-i Apollo-Picarro, USA) to quantify both the DIC concentration and the carbon isotope ratio (δ^13^C) at initial time.

After quantifying the initial DIC, the reaction was initiated by adding an equimolar amount of RuBP. The syringe was incubated on a roller at 25 °C, and aliquots were withdrawn over a period of 6 to 8 h until around 70-80% of the initial DIC was consumed. Each aliquot was processed identically to the initial sample, and DIC concentration and δ^13^C values were measured throughout the course of the reaction using the Apollo-Picarro system.

Additional 0.1 to 0.5 mL samples were transferred to 5 mL septum-capped vials containing 200 mM HPO_3_ flushed with helium. These were analyzed using a GasBench II connected to an isotope ratio mass spectrometer (Delta V Plus, Thermo Fisher Scientific, USA), serving to validate the isotope ratios obtained with the Apollo-Picarro system.

ε_Rubisco_ was calculated using the linearized Rayleigh equation, as described by (Scott et al., 2004):

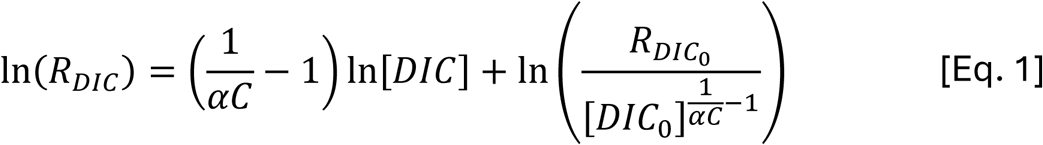

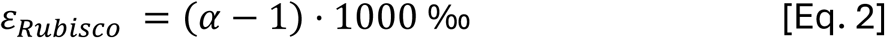

Where R_DIC_ is the ^13^C/^12^C ratio of the residual DIC, [DIC] is the concentration of dissolved inorganic carbon, R_DICO_ and [DIC_0_] are their corresponding initial values, C is the equilibrium isotope effect between CO_2_ and HCO_3_^-^, and α is the kinetic isotope effect. Experiments were performed in triplicate, and ε_Rubisco_ results were combined using the Pitman estimator (Scott et al., 2004). Uncertainties are reported as the standard error and the corresponding 95% confidence intervals (Supplementary Table 1). *Synechococcus* sp. PCC 6301 and *Spinacia oleracea* ε_Rubisco_ values were obtained from Wijker et al. (2025).

## Modelling Rubisco Assimilation

Farquhar’s biochemical model of photosynthesis (Farquhar et al., 1980) was adapted to estimate Rubisco-limited assimilation rate per μmol of active sites (A_Rub_) at 25 °C using the kinetic data obtained for each cyanobacterial strain in this study. For comparative purposes, A_Rub_ of the C_3_ crop model *Triticum aestivum* was also calculated using Rubisco kinetic data measured in the present work at 25 °C (Table 1). The equation used was:

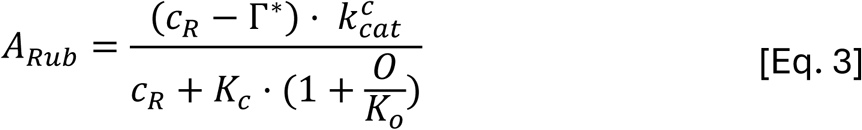

Where 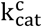 is the carboxylation turnover rate, K_c_ and K_o_ the Michaelis-Menten semi-saturation constants for the CO_2_ and O_2_, respectively. C_R_ and O are the partial pressures of CO_2_ and O_2_ at the Rubisco catalytic sites. 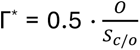, where S_c/o_ is Rubisco specificity factor measured at 25 °C. Henry’s law was used to convert liquid-phase kinetic values to gas-phase, using a CO_2_ solubility constant of 0.0328 mol L^-1^ Bar^-1^ and a O_2_ solubility constant of 0.00119 mol L^-1^ Bar^-1^ (Galmés et al., 2016).

The percentage of maximum Rubisco assimilation at an ambient CO_2_ concentration of 400 ppm (A_Rub max_ at C_a_ = 400 ppm) was calculated by multiplying the CCM effectiveness of each species by C_a_ of 400 ppm. The resulting CO_2_ concentration at the vicinity of Rubisco catalytic sites (C_R_) and the gas-phase Rubisco kinetic parameters were applied to the A_Rub_ model [Eq. 3] to calculate Rubisco assimilation at C_a_ = 400 ppm (A_Rub_ at C_a_ = 400 ppm). Then A_Rub max_ at C_a_ = 400 ppm was calculated by dividing A_Rub_ at C_a_ = 400 ppm by A_Rub max_ (A_Rub_ when C_R_ tended to infinite) and multiplied by 100.

## Phylogenetic Analyses

Rubisco large- and small-subunit gene sequences (*rbcL* and *rbcS*, respectively), available in GeneBank, were retrieved for the 12 species of Cyanobacteria with published Rubisco kinetic data, summarized in Supplementary Spreadsheet 1 (accession numbers and references in Supplementary Table 2). Sequences were aligned using MUSCLE algorithm and manually trimmed. A phylogenetic tree was constructed (Supplementary Figure 1) using maximum likelihood method with 1,000 bootstrap replicates, applying the Tamura–Nei substitution model with Gamma distribution and a proportion of invariant sites (Nei and Kumar, 2000; Tamura et al., 2011). All analyses were performed in MEGA 12 (version 12.0.11). Phylogenetic signal of Rubisco kinetic parameters for the 12 cyanobacterial species was assessed by calculating Bolmberg’s K (Supplementary Table 3) using the *picante* package (version 1.8.2) in R (Blomberg et al., 2003).

## Statistical Analyses

Normality and homoscedasticity of variances were assessed using Anderson–Darling and Levene’s tests, respectively. For parametric data, differences among group means were tested using one-way analysis of variance (ANOVA), followed by Tukey’s post hoc test for pairwise comparisons. For non-parametric data, the Kruskal–Wallis test was used, followed by Dunn’s post hoc test with Bonferroni adjustment. Differences between two groups were assessed using Student’s t-test. For ε_Rubisco_, Pitman estimator means were calculated in MATLAB (version R2024b) and pairwise comparisons were performed using z-test with Holm correction for multiple testing. A p-value below 0.05 was considered statistically significant for all tests. All statistical analyses were conducted in R (version 4.3.1) and RStudio (version 2023.06.1). Data visualization was performed using the ggplot2 package (version 3.5.2).

## Results

### Rubisco Kinetic Parameters are Diverse Across Cyanobacterial Lineages

Phylogeny (Supplementary Figure 1) explained little of the kinetic variation with Blomberg’s K values below 1 indicating weak phylogenetic signal (Supplementary Table 3) (Blomberg et al., 2003). Nevertheless, a large variability was observed in the Rubisco kinetic parameters among cyanobacterial lineages.

Rubisco specificity factor (S_c/o_) values varied among the species analyzed, ranging from the highest in *G. violaceus* and *C. thermalis* (74.6 and 66.0 mol mol^-1^, respectively) to the lowest in *Synechococcus* sp. (48.2 mol mol^-1^; Table 1). The highest K_c_ value was measured in *O. acuminata* (180.8 μM), whereas the lowest values were observed in *C. thermalis* and *C. aponinum* (87.3 and 80.0 μM, respectively). 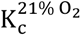 was approximately two-fold higher in *Synechococcus* sp. And *O. acuminata* (224.0 and 214.3 μM, respectively) than in *C. aponinum* and *C. thermalis* (Table 1). The K_o_ values of *C. aponinum* and *Synechococcus* sp. were about half those of the other species. The highest 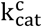 was recorded in *G. violaceus* (11.8 s^-1^), although not significantly different from the values in *C. thermalis* and *O. acuminata*. The 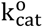 of *C. thermalis* was more than two-fold higher than that of *Synechococcus* sp. (Table 1). Carboxylation efficiencies 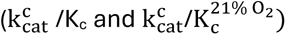 were around two-fold higher in *C. thermalis* and *C. aponinum* compared with *O. acuminata* and *Synechococcus* sp., with similar patterns observed for oxygenation efficiency 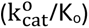.

*Compared to the determined Synechococcus* sp. ε_Rubisco_ (22.7‰) with this method (Wijker et al., 2025), other cyanobacterial species had higher ε_Rubisco_ with *G. violaceus* and C. *thermalis* exhibiting the highest values (27.3 and 26.1‰, respectively; Table 1). The percentage of Rubisco relative to the total soluble protein (% Rubisco to TSP) ranged from 0.8% in *Synechococcus* sp. to 1.4% in *G. violaceus* and *C. thermalis* (Table 1).

When compared with previously published data (Figure 1 and Supplementary Spreadsheet 1), the results of the present study stand out for exhibiting the highest or lowest values in most Rubisco kinetic parameters in Cyanobacteria. *C. aponinum* and *C. thermalis* from present study exhibited the lowest K_c_ values ever measured, similar to those of *Aphanocapsa virescens* (K_c_ = 80 μM; Jordan et al., 1983). In contrast, α-cyanobacterium *Prochlorococcus marinus* MIT 9313 showed the highest K_c_ value reported so far (Shih et al., 2016; Figure 1A). *Gloeobacter violaceus* 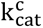 value (present study) was the highest reported for Cyanobacteria (Figure 1B). The K_o_ value of *O. acuminata* represented the highest value in the phylum, while that of *C. aponinum* was the lowest (both from the present study, Figure 1C). The highest 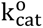 was observed in *C. thermalis* (present study), whereas the lowest was reported in *Anabaena* sp. M-3 by Badger (1980; Figure 1D). S_c/o_ across Cyanobacteria ranged from 32.0 mol mol^-1^ in *Leptolyngbya boryana* reported by Kent and Tomany (1984) to 74.6 mol mol^-1^ in *G. violaceus* (present study). Reported ε_Rubisco_ values varied from 22.7‰ in *Synechococcus* sp. PCC6301 to 27.3‰ in *G. violaceus* (Figure 1F). The highest value of the Rubisco catalytic carboxylation efficiency 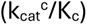 was obtained in *C. thermalis* (present study) and the lowest value in *P. marinus* (in Shih et al., 2016; Figure 1G), while the Rubisco catalytic oxygenation efficiency 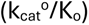 ranged from 0.2 s^-1^ mM^-1^ in *Anabaena* sp. (in Badger, 1980) to 1.7 s^-1^ mM^-1^ in *C. aponinum* (present study).

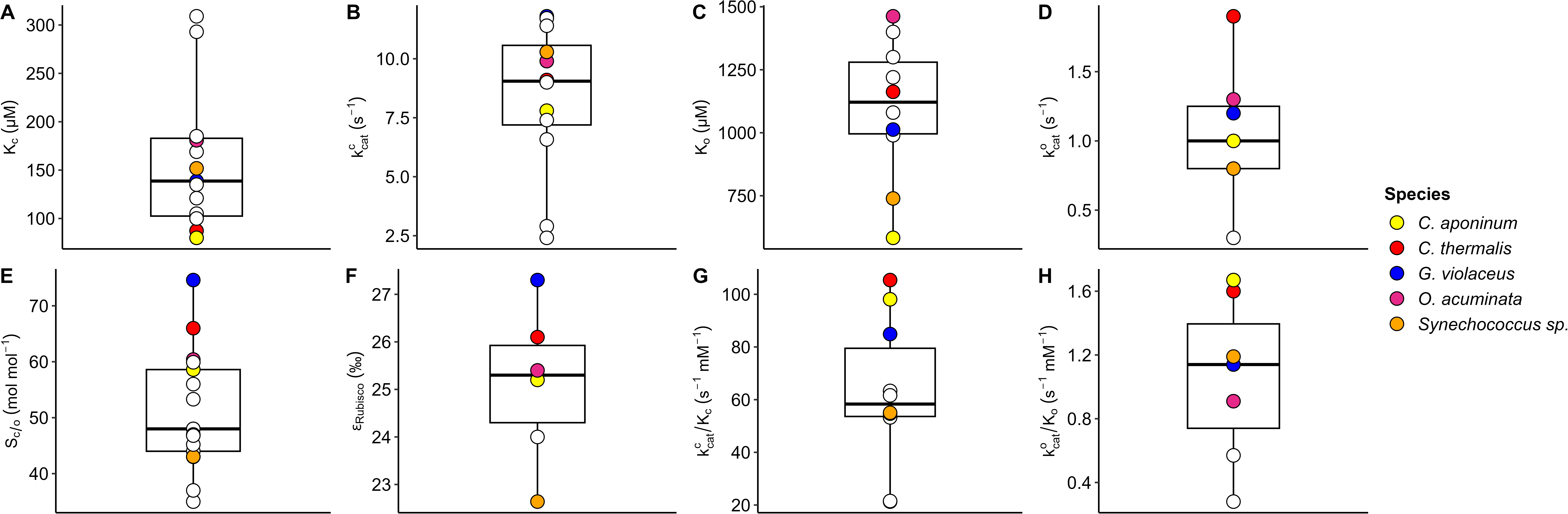
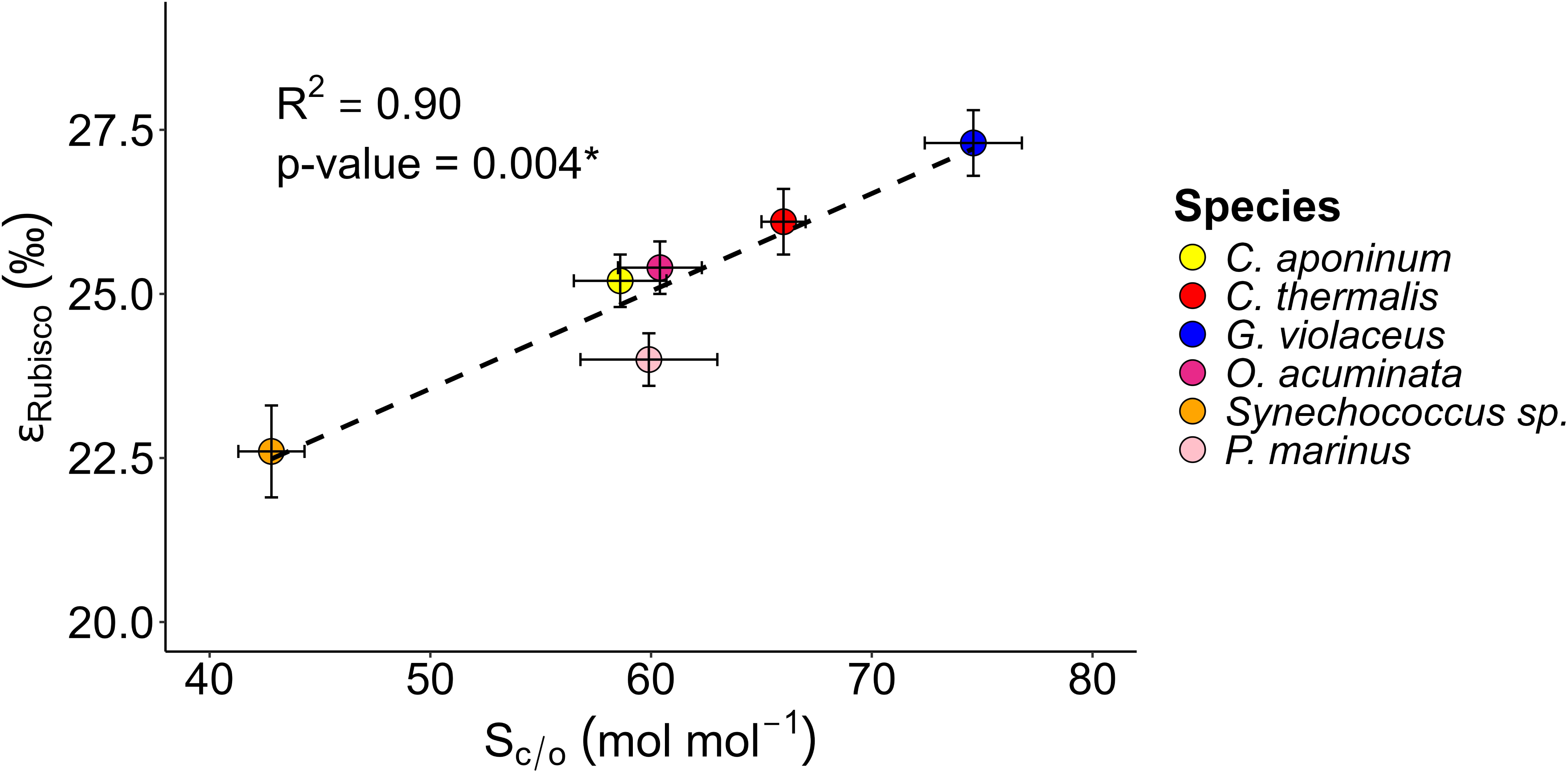
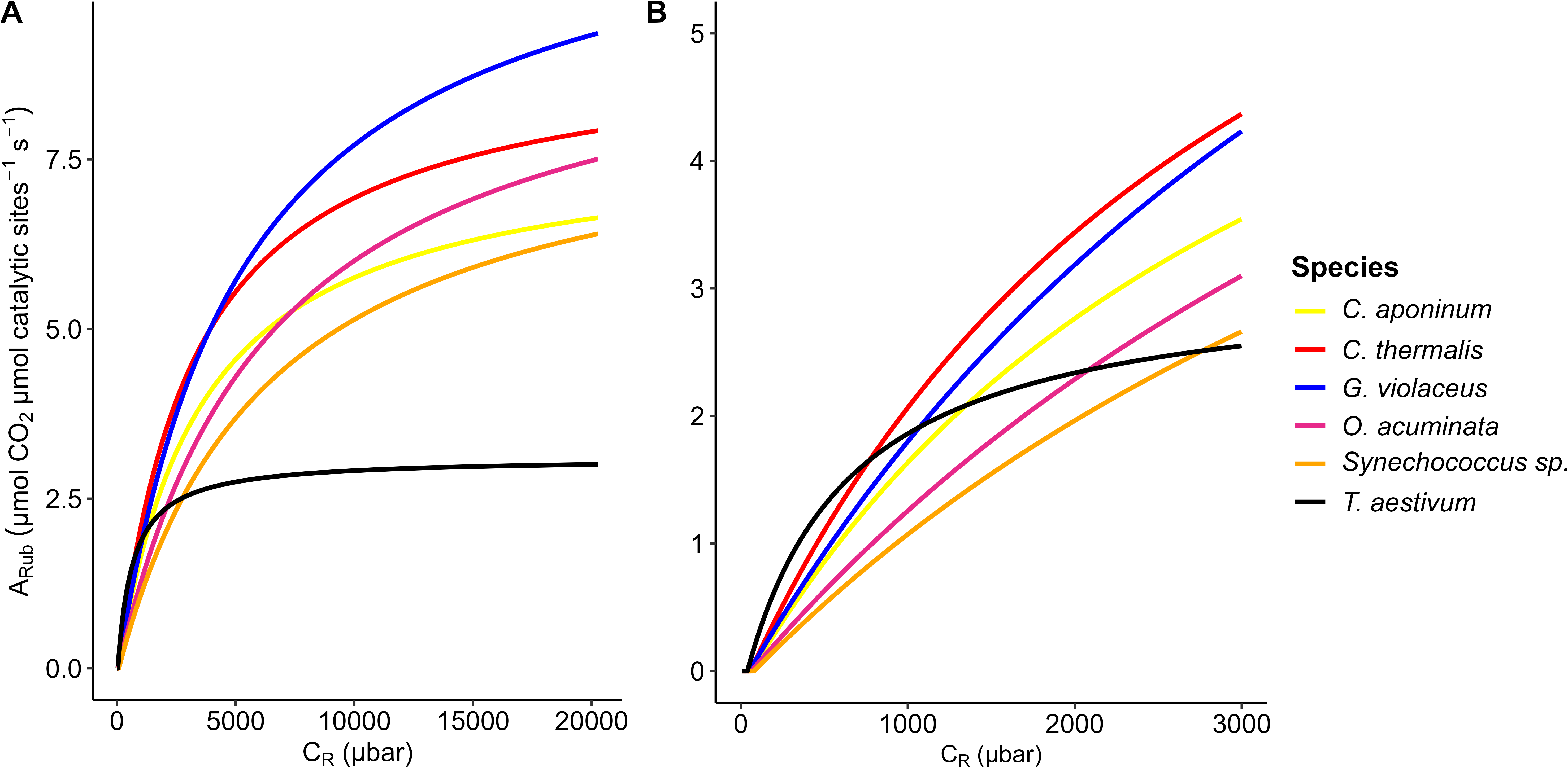

There was a positive correlation between ε_Rubisco_ and S_c/o_ (p-value = 0.004, R^2^ = 0.90) including *P. marinus*, for which ε_Rubisco_ values from Scott et al. (2007) and S_c/o_ from Shih et al. (2016) were incorporated (Figure 2). The classical trade-offs between Rubisco kinetic parameters 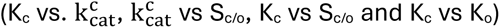 were not found in Cyanobacteria (Supplementary Figure 2).

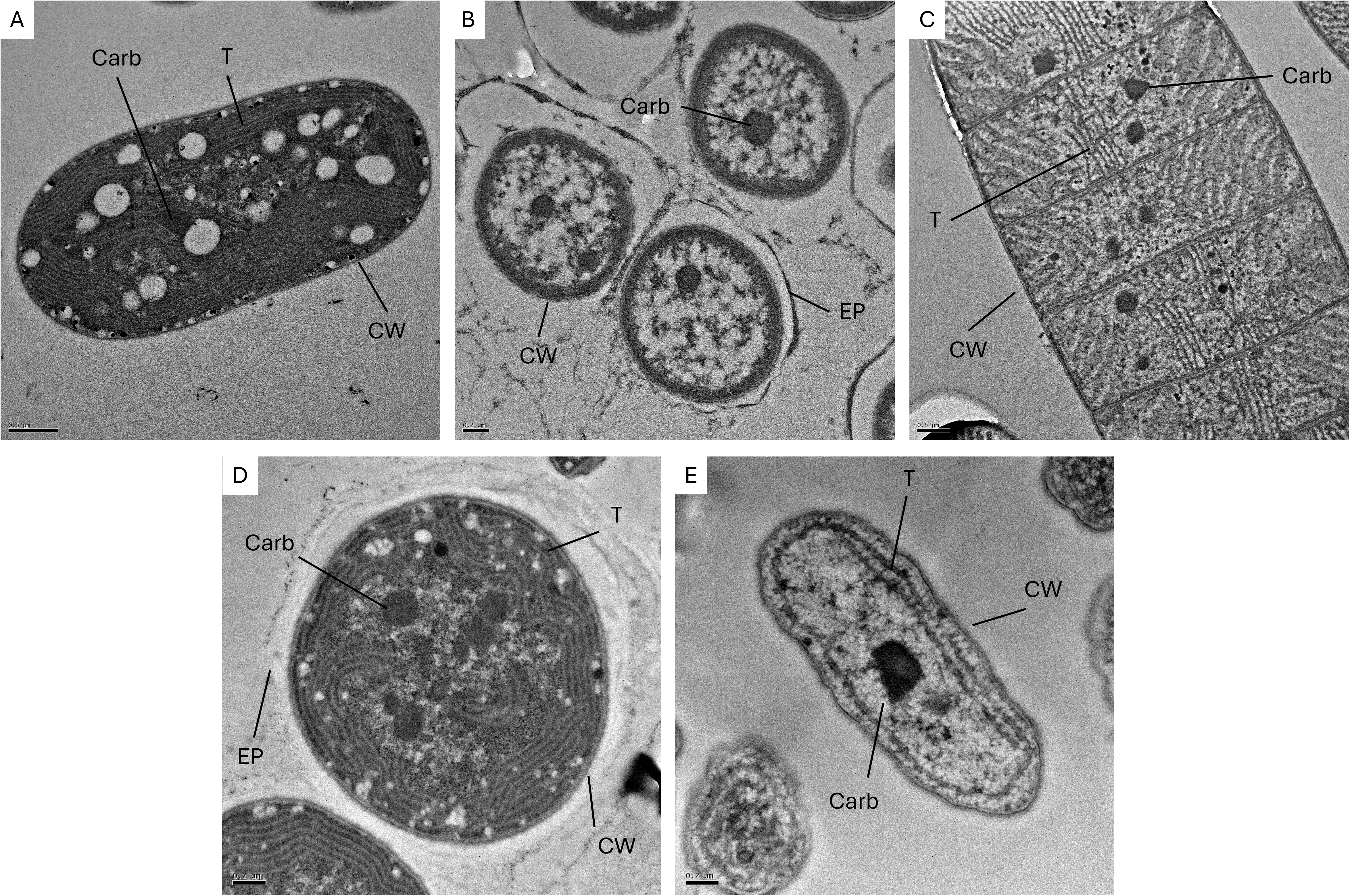

Among all studied strains, *Gloeobacter violaceus* presented the highest A_Rub_ values at C_R_ greater than 4,000 ppm (Figure 3A). All cyanobacterial strains presented a lower Rubisco assimilation rate (A_Rub_) than the C_3_ model plant *T. aestivum* across the range of C_R_ from 0 to 770 ppm (Figure 3B). Among the strains, *C. thermalis* intersected *T. aestivum* A_Rub_ at the lowest C_R_, and *Synechococcus* sp. at the highest.

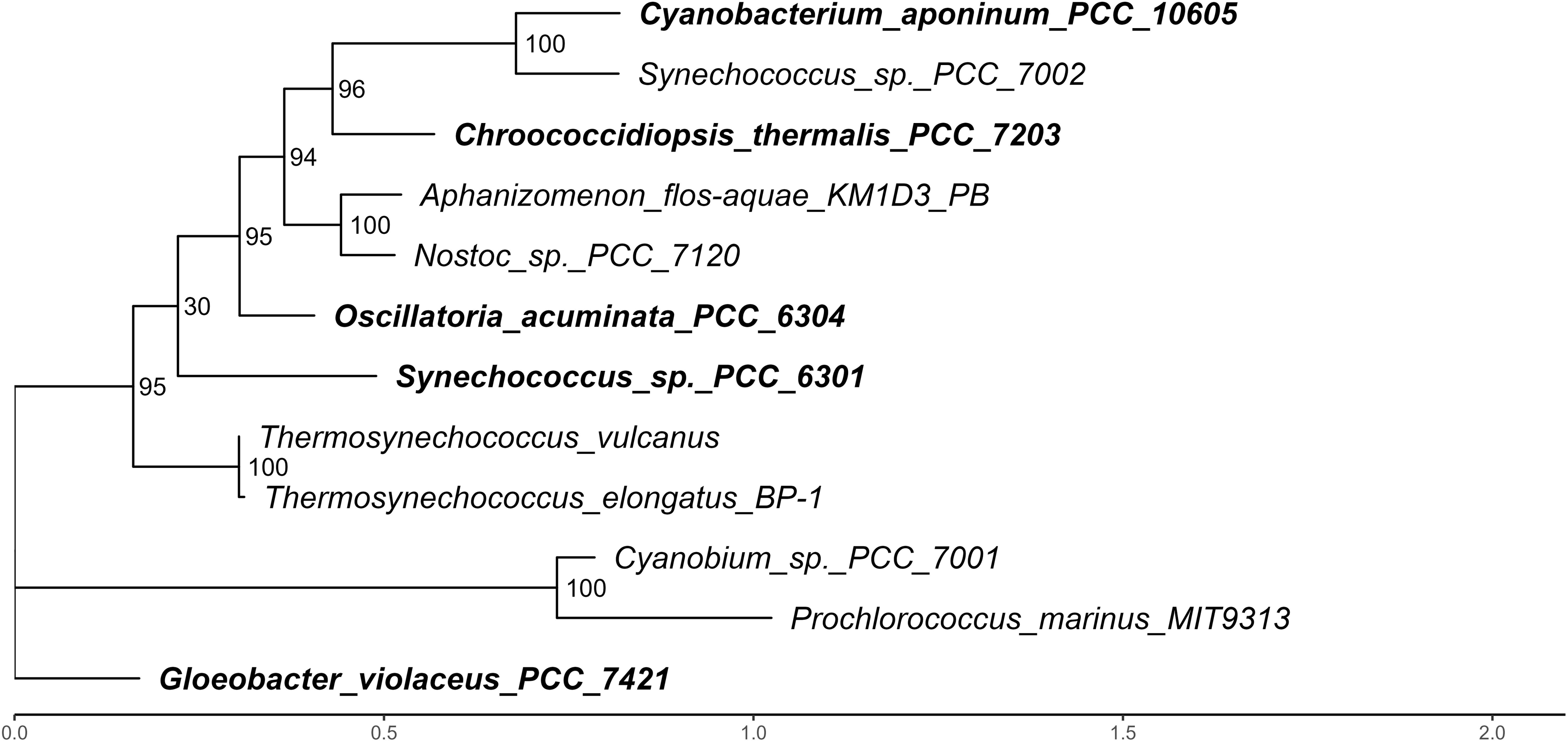
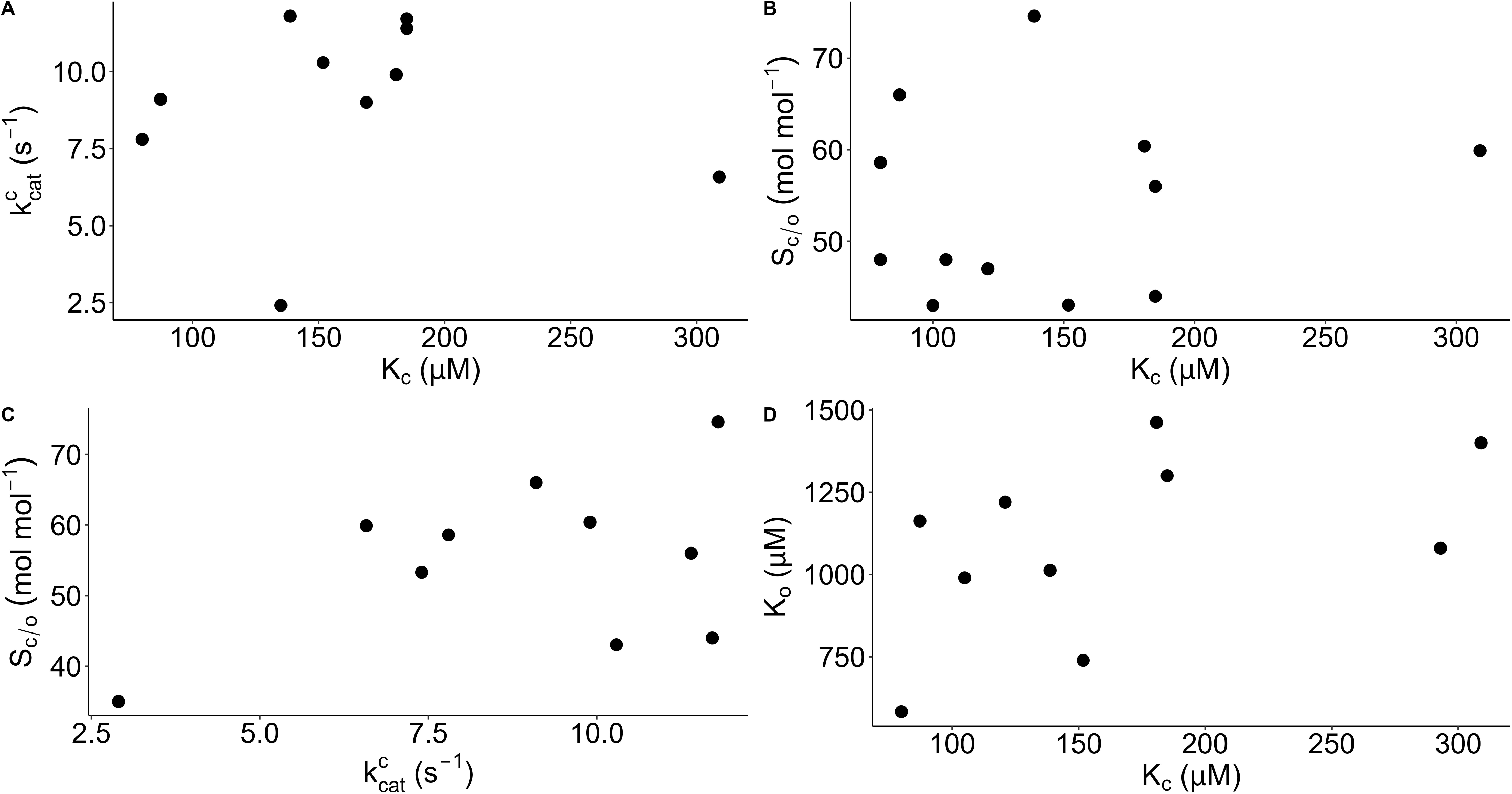

### Variability of CO_2_-Concentrating Mechanisms across Phylogenetically Distant Cyanobacteria

The presence of carboxysomes was confirmed in all examined strains through TEM imaging (Figure 4). Different cell morphologies were observed among the different species: *Oscillatoria acuminata* formed long filamentous strips, while *C. thermalis* and *G. violaceus* formed tetrads enclosed in exopolysaccharide matrices. In contrast, *Synechococcus* sp. and *C. aponinum* appeared free-living coccoid cells. In addition, *C. aponinum*, *C. thermalis*, and *O. acuminata* presented an extensive network of internal thylakoid membranes, whereas *Synechococcus* sp. showed fewer internal membranes, and *G. violaceus* completely lacked thylakoids, consistent with previous observations by Rippka et al. (1974) (Figure 4).

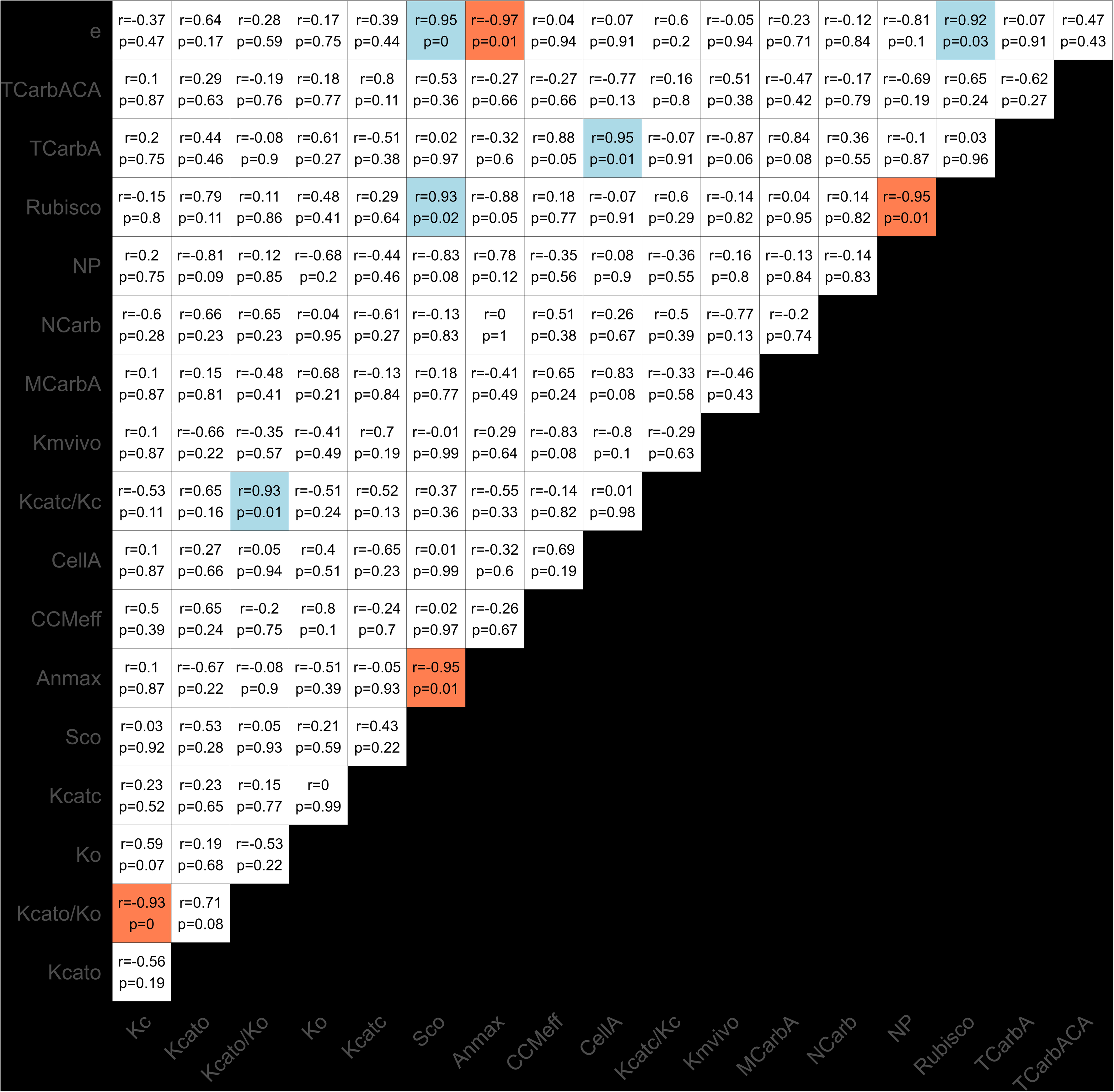

The largest average cell area (CellA) was recorded for *O. acuminata* (6.3 μm^2^) and *C. aponinum* (5.0 μm^2^), whereas the smallest was observed in *G. violaceus* and *Synechococcus* sp. (1.2 and 1.5 μm^2^, respectively; Table 2). The highest average number of carboxysomes per cell (NCarb) was found in *C. thermalis* (2.8 per cell), while *G. violaceus* had the lowest (1.1 per cell; Table 2). *Oscillatoria acuminata* displayed both the largest mean carboxysome area (MCarbA = 76 nm^2^) and the greatest total carboxysome area per cell (TCarbA = 112 nm^2^). The highest proportion of cell area occupied by carboxysomes (TCarbA/CellA) was observed in *G. violaceus* and *C. thermalis* (2.7 and 2.3%, respectively; Table 2).

Regarding the *in vivo* photosynthetic measurements (Table 3), extracellular CAs inhibitor, AZ, had a significant effect on A_n_ only in *Synechococcus* sp. (11.6% net photosynthetic inhibition—NPI) and *C. aponinum* (4.3% NPI). In contrast, the internal and external CAs inhibitor, EZ, significantly reduced A_n_ in all strains, with a NPI ranging from 10.2% in *C. aponinum* to 43.4% in *Synechococcus* sp. (Table 3). Similarly, the anion-exchange inhibitor DIDS caused significant NPI in all strains, with the greatest effect observed in *C. thermalis* (22.0%), threefold higher than the observed for *O. acuminata* (6.7%; Table 3).

*In vivo* measurements of the net photosynthetic rate under different DIC concentrations (photosynthesis-C curves) revealed that *Synechococcus* sp. exhibited a maximum assimilation rate (A_n max_) nearly seven-fold higher than that of *G. violaceus*. In contrast, the other species displayed similar A_n max_ values (Table 4).

The lowest *in vivo* Michaelis–Menten semi-saturation constant for CO_2_ (K_m_ *_in vivo_*) was observed in *C. thermalis* (0.5 µM), five-fold lower than that of the highest value observed in *G. violaceus* (2.7 µM; Table 4). However, all K_m *in vivo*_ values measured in the different cyanobacterial strains indicate that net photosynthesis must be almost saturated at the environmental CO_2_ levels.

### Evolutionary Interplay Between CO_2_-Concentrating Mechanisms and Rubisco in Distantly Related Cyanobacteria

The ratio between 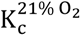 and K_m *in vivo*_ is an indicative of the strength of the CCMs in elevating CO_2_ levels around Rubisco active sites. It also acts as a proxy for estimating how many times ambient CO_2_ is concentrated inside the cell. According to Raven et al. (2017), a 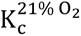: K_m_ *_in vivo_* ratio greater than 2.5 indicates the presence of effective CCMs. All cyanobacterial strains examined here exhibited strong CCMs, with *O. acuminata* reaching internal CO_2_ concentrations up to 243.8 times higher than ambient levels—approximately four times greater than those observed in *G. violaceus* (Table 4).

The highest percentage of A_Rub max_ at C_a_ = 400 ppm (Supplementary Table 4) was observed in *C. thermalis* (96.4 %), whereas *G. violaceus* showed the lowest values (83.3 %). However, all species present Rubisco saturation values higher than 80 %, indicating that their CCMs are highly effective.

## Discussion

### Lineage-Specific Features of Cyanobacterial CO_2_-Concentrating Mechanisms

All cyanobacterial strains analyzed exhibited exceptionally high *in vivo* CO_2_ affinity (K *_in vivo_* < 3 μM; Table 4), surpassing values reported for other CCM-expressing photoautotrophs (e.g. pyrenoid-containing green algae; Capó-Bauçà et al., 2024). The high values of CCM effectiveness (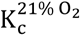: K_m *in vivo*_ ratio) also confirm the elevated capacity to concentrate CO_2_ near Rubisco active sites, which has been previously associated to the presence of carboxysomes in Bacteria (Raven et al., 2017).

Among the species analyzed, *C. thermalis* and *O. acuminata* exhibited the lowest K_m *in vivo*_ and the most effective CCMs (Table 4). In *C. thermalis*, these traits can be related to its high number of carboxysomes per cell (Table 2), allowing Rubisco to operate close to CO_2_-saturation (Supplementary Table 4). These features likely represent adaptations to hot, arid environments where CO_2_ solubility is inherently low (Skirrow, 1975). In the case of *O. acuminata*, efficient CCMs may stem from its large carboxysome area, despite a relatively low carboxysome-to-cell area ratio (Tables 2). Its filamentous multicellularity—an early cyanobacterial innovation preceding eukaryotes by hundreds of millions of years—may also confer advantages such as reduced CO_2_ leakage and more efficient resource allocation.

Conversely, the relatively lower CCM effectiveness found in *G. violaceus* (Table 4) likely reflects its minimal CCM architecture, with the smallest reported β-carboxysome gene content, including only two CO_2_ uptake systems and a single bicarbonate transporter and CA located at the plasma membrane (Long et al., 2016). This lineage diverged ∼2.8 billion years ago, when atmospheric CO_2_ vastly exceeded O_2_ concentrations. That imposed little selective pressures to evolve a more complex CCM (Raven and Sánchez-Baracaldo, 2021). These ancestral features together with lack of thylakoids may limit photosynthetic efficiency resulting in slow growth (Raven and Sánchez-Baracaldo, 2021). These traits may have persisted in the extant strain likely due to hypobradytely—slow evolutionary rate (Schopf, 1994).

Inhibitor assays showed that AZ (external CA inhibitor) had little effect on net photosynthesis, while EZ (both external and internal CAs inhibitor) strongly inhibited it in all strains (Table 3). This highlights the important role of internal CAs in the CCMs of the β-cyanobacteria explored here. Concretely, β-cyanobacteria CCMs consist in up to three bicarbonate transporters (BicA, SbtA, and BCT1) and up to two CO_2_ uptake systems (NDH-1_3_ and NDH-1_4_), these last having CA activity and being sensitive to EZ inhibition, as confirmed in *Synechocystis* sp. PCC 6803 and Synechococcus sp. PCC 7942 (Zhang et al., 2025). In addition, β-cyanobacteria present β-carboxysomes—bigger and more complex than α-carboxysomes. β-carboxysomes consist in a proteinic shell of BCM protein forming flat hexagonal tiles with small pores. In the vertices where five sheets meet, there is a CcmL protein as a pentagonal pyramid (Espie and Kimber, 2011). Inside the shell, the CA CcmM binds Rubisco and other proteins to form a condensed aggregation (Espie and Kimber, 2011). Internal CAs of β-carboxysomes (CcmM and CcaA) are also highly sensitive to EZ (Peña et al., 2010). DIDS also reduced photosynthesis, consistent with reliance on active HCO_3_ ^-^ uptake by plasma-membrane anion-exchange transporters rather than extracellular CAs activity (Long et al., 2016). Distinct species-specific responses to inhibitors suggest lineage-specific configurations of CCM components, probably expressing differential CCM components in response to adaptation to each environmental condition. The efficiency of cyanobacterial CCMs enables cells to allocate less nitrogen to Rubisco compared to autotrophs that lack such mechanisms (Table 1). This is consistent with the low Rubisco levels reported in other CCM-expressing aquatic phototrophs such as diatoms (Young et al., 2016).

### Rubisco Kinetic Diversity across Cyanobacterial Lineages and Co-evolution with CCMs

Cyanobacterial lineages display diverse Rubisco kinetics despite a shared evolutionary signature of high 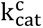 and K_c_ (therefore, low affinity for CO_2_), shaped by co-trait variation with highly effective CCMs. Among the strains analyzed and compiled, the highest Rubisco CO_2_ affinities found in *C. thermalis* and *C. aponinum* might be a consequence of selective pressure under limited CO_2_ environments dominated by high temperatures (Tang et al., 2022). Concretely, *C. thermalis* inhabits desertic areas and *C. aponinum* some thermal springs, with both being thermotolerant species (Chen et al., 2025). These findings support the idea that some of the most distinctive Rubisco kinetic parameters are found in extremophiles (Aguiló-Nicolau et al., 2024). In addition, both species exhibiting the highest carboxylation efficiencies (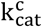/K_c_) indicates that not only its Rubisco presents high affinity for CO_2_ but also that it has co-evolved with highly efficient CCMs, forcing the evolution towards high 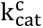, as they are able to concentrate carbon more than 200 times C_a_ at the vicinity of Rubisco (e.g. *C. thermalis*; Table 4), achieving near-saturation Rubisco microenvironment (Supplementary Table 4).

By contrast, α-cyanobacterium *P. marinus* exhibiting the lowest CO_2_ affinity while presenting relatively high 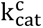 (Shih et al., 2016) may reflect its Rubisco co-evolution with streamlined but efficient CCMs, which comprise one or two low-affinity, high-flux bicarbonate transporters and carboxysomes, lacking CO_2_ uptake systems (Hopkinson et al., 2014). This configuration results in elevated C_R_, relaxing selection for high affinity enforcing faster turnover.

*Gloeobacter violaceus* high S_c/o_ may be the result of its co-evolution with relatively inefficient CCMs. However, as shown in Supplementary Table 4, its assimilation rate at C_a_ = 400 ppm is already close to its maximum—as well as in all other species—indicating that CCMs can elevate CO_2_ concentrations sufficiently to almost saturate Rubisco. Interestingly, *G. violaceus* also showed the highest 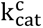 within the phylum, contrasting with previously reported values (0.016 s^-1^) from heterologous expression in *E. coli* construct (Sidhu et al., 2021)—likely underestimated due to non-saturating substrate conditions, over-estimated protein quantification or enzyme deactivation. Our measured value aligns with other Cyanobacteria (see Supplementary Spreadsheet 1) and probably consistent with its close to Rubisco saturation microenvironment (Supplementary Table 4).

Reconstructions of ancestral cyanobacterial Rubiscos suggest that extant β-cyanobacteria retain similar S_c/o_ and K_c_ but double the 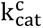 of ancestral form IB Rubisco (Shih et al., 2016). Consistent with this pattern, the K_c_ value of *G. violaceus* aligns with evolution under high-CO_2_ conditions during Archean-early Proterozoic. After the emergence of CCMs, probably during the GOE (Hurley et al., 2021), although still under debate, selection appears to have increased 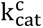 while leaving K_c_ largely unchanged. A shift toward higher S_c/o_ may also have occurred in *G. violaceus*, potentially reflecting its less effective CCM relative to other species. Likewise, *C. thermalis* and *C. aponinum* show higher S_c/o_ and 1/K_c_ than the reconstructed ancestral IB enzyme (Shih et al., 2016). Together, these trends—higher 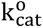 associated with increasingly efficient CCMs and relative conservation of K_c_—yield greater carboxylation efficiency in modern cyanobacterial Rubiscos.

While the relative importance of kinetic trade-offs versus phylogenetic constraints in Rubisco evolution remains debated (Bouvier et al., 2021; Tcherkez and Farquhar, 2021), our results suggest that neither dominates in Cyanobacteria. Notably, we did not observe the trade-offs among Rubisco kinetic parameters that are well-documented in higher plants—an absence previously observed in diatoms and microalgae (Young et al., 2016) (Supplementary Figure 2).

### Rubisco Trade-of Between S_c/o_ and ε_Rubisco_ in Cyanobacteria

The observed positive correlation between S_c/o_ and ε_Rubisco_ in our dataset is consistent with the original prediction that a more product-like carboxylation transition state increases intrinsic isotope discrimination (Tcherkez et al., 2006). A recent meta-analysis reported substantial scatter in this relationship when data from different photosynthetic groups were pooled, in part because paired ε_Rubisco_-S_c/o_ measurements are often missing (Wang et al., 2023a). By providing both parameters for multiple cyanobacterial lineages, our results indicate that, within this phylum, a common mechanism regulates variation in S_c/o_ which may allow estimation of ε_Rubisco_ from kinetic data or vice versa.

Whether ancestral enzymes displayed higher or lower ε_Rubisco_ discrimination is still debated (Kędzior et al., 2022; Wang et al., 2023b). Our finding that the highest ε_Rubisco_ occurs in the basal cyanobacterial strain *G. violaceus* is consistent with a model of gradually declining discrimination through geological time. This pattern aligns with reported decreases in isotopic discrimination of plant biomass (Kohn, 2010) and with ancestral Rubisco reconstructions in *S. elongatus* (Kędzior et al., 2022). However, alternative analyses argue for relatively constant ε_Rubisco_ over Earth history (Garcia et al., 2021). Resolving this discrepancy will require broader phylogenetic sampling of paired ε_Rubisco_ and S_c/o_, including potentially resurrected ancestral enzymes as demonstrated by Kędzior et al. (2022). However, Rubisco kinetics measured from heterologously expressed enzymes should be interpreted with caution, as they may not fully reflect native behavior.

The carbon-isotope record in sedimentary rocks has long been used to infer ancient atmospheric CO_2,_ assuming that *in vivo* fractionation preserved in the fossil biomass responds dominantly to variation in the degree of limitation of [CO_2_]_aq_. Physical models for microalgae assuming a constant ε_Rubisco_ (Rau et al., 1996) interpret declines in the photosynthetic fractionation (ε_p_) between biomass and the [CO_2_]_aq_ source as evidence for declining CO_2_. Our finding of highest ε_Rubisco_ in the basal strain compared to *Synecococcus* may indicate a progressive 5‰ decrease in ε_Rubisco_ over the late Proterozoic-early Paleozoic (∼0.8-0.35 Ga) concomitant with the evolution of CCM with declining atmospheric CO_2_. If similar ε_Rubisco_ declines characterized other phytoplankton, it implies that a portion of the observed long term (ε_p_) decline (Witkowski et al., 2018) may be attributed to the evolution of Rubisco with declining CO_2_ rather than changes in cellular C fluxes with progressive C limitation. At the same time, the significance ε_Rubisco_ relative to fractionation by other enzymes contributing to the CCM of modern microalgae remains under discussion (Wilkes and Pearson, 2019).

Beyond paleo-inference, mapping ε_Rubisco_ diversity in Cyanobacteria has immediate implications. As major contributors to Precambrian sedimentary records, lineage-resolved ε_Rubisco_ can help constrain the timing of photosynthetic evolutionary milestones (e.g. PSII emergence) when integrated with geochemical archives. Mechanistically, the ε_Rubisco_-S_c/o_ trade-of informs how transition-state structure shapes Rubisco catalysis and can guide selection or engineering of variants with desirable kinetic properties.

## Concluding remarks

This study reveals substantial variation in Rubisco kinetics and CCMs across phylogenetically and ecologically diverse cyanobacterial lineages, yet a shared evolutionary trend of high catalytic turnover and low CO_2_ affinity, consistent with a coevolution with highly effective CCMs. Notably, extremotolerant strains possessed Rubiscos with enhanced CO_2_ affinities and stronger CCMs, while the basal lineage displayed a minimal CCM, and high CO_2_/O_2_ specificity, catalytic turnover and ε_Rubisco_, likely reflecting Rubisco co-evolution with these minimal CCMs.

We identified a robust positive correlation between S_c/o_ and ε_Rubisco_, providing empirical support for theoretical predictions linking enzyme CO_2_ specificity and C isotope effects. This relationship suggests that ε_Rubisco_ may serve as a practical proxy for S_c/o_ an vice versa within Cyanobacteria and has important implications for interpreting stable isotope signatures in both modern ecosystems and deep-time records.

Altogether, our findings underscore the dynamic interplay between Rubisco evolution and CCMs in Cyanobacteria, highlighting the importance of the intracellular CO_2_/O_2_ ratio as a dominant driver of Rubisco evolution rather than phylogenetic constraints or inherent kinetic trade-offs. Moreover, this work emphasizes the need to expand biochemical and isotopic datasets beyond model organisms to fully capture the enzymatic diversity across the phylum. Such knowledge not only advances our understanding of cyanobacterial evolution and ecology but also sets a solid basis to efforts aimed at enhancing carbon fixation in crops by integrating cyanobacterial components.

## Acknowledgements

This work was financially supported by the Spanish Ministry of Sciences, Innovation and Universities, the Spanish State Research Agency and the European Regional Development Funds (project UNRAVENAR, PID2023-148523NB-I00) funded to Jeroni Galmés. Pere Aguiló Nicolau was supported by a pre-doctoral grant from the Government of the Balearic Islands (FPI/046/2020). Technical instrumentation was funded by the platform HiTech-INAGEA (SINCO 2022/18198) with support from the Conselleria d’Educació i Universitats, Govern de les Illes Balears, and FEDER 2021-2027. Rubisco fractionation measurements were supported by Grant ETH-03 19-1 to Heather Stoll. We thank Trinidad Garcia for technical help and organization of the radioisotope facilities at the Serveis Cientifico-Tècnics (UIB), Madalina Jaggi and Stewart E. Bishop for their technical support at ETH-Zürich and Toni Palerm for his advice in statistical analysis. We used ChatGPT (OpenAI) to improve wording and to draft initial R code for figure generation. All text and code were reviewed, edited and validated by authors, who take full responsibility for the content.

## Author Contributions

JG and HS conferred the idea of the manuscript with help of CI and SCB. PAN, CI, SCB and RW developed the experimental design. PAN and RW executed the experimental design. PAN analyzed the data, generated the tables and figures, and wrote the manuscript body. All coauthors reviewed the manuscript thoroughly.

