## Supplementary material for "Diversity in Rubisco Kinetics and CO₂-Concentrating Mechanisms Among Cyanobacterial Lineages": Figures and tables file

**Table 1.** *In vitro* Rubisco kinetic parameters at 25 °C among the different cyanobacterial strains analysed (mean ± standard error).

| **Species** | **S_c/o_ (mol·mol^-1^)** | **K_c_ (μM)** | $\mathbf{K}_{\mathbf{c}}^{\boldsymbol{21 \%}\mathbf{O}_{\mathbf{2}}}$ **(μM)** | **K_o_ (μM)** | $\mathbf{k}_{\mathbf{cat}}^{\mathbf{c}}$ **(s-1)** | $\mathbf{k}_{\mathbf{cat}}^{\mathbf{o}}$ **(s-1)** | $\mathbf{k}_{\mathbf{cat}}^{\mathbf{c}}$ **/K_c_**  **(s^-1^ mM^-1^)** | $\mathbf{k}_{\mathbf{cat}}^{\mathbf{c}}$**/**$\mathbf{K}_{\mathbf{c}}^{\boldsymbol{21\%}\mathbf{O}_{\mathbf{2}}}$ **(s^-1^ mM^-1^)** | $\mathbf{k}_{\mathbf{cat}}^{\mathbf{o}}$**/K_o_**  **(s^-1^ mM^-1^)** | **% Rubisco to TSP** | ε**_Rubisco_ (‰)** |
| --- | --- | --- | --- | --- | --- | --- | --- | --- | --- | --- | --- |
| *C. aponinum* | 58.6 ± 2.1 b | 80.0 ± 1.3 c | 116.1 ± 3.8 c | 582.9 ± 45.9 b | 7.8 ± 0.1 b | 1.0 ± 0.1 bc | 98.1 ±s 1.3 a | 67.7 ± 2.1 b | 1.67 ± 0.02 a | 1.0 ± 0.1 ab | 25.2 ± 0.4 b |
| *C. thermalis* | *66.0 ± 1* ab | *87.3 ± 0.8* c | *105.4 ± 2.4* c | *1162.6 ± 105.3* a | *9.1 ± 0.2* ab | *1.9 ± 0.1* a | *105.4 ± 1.5* a | *86.0 ± 0.8* a | *1.60 ± 0.02* a | 1.4 ± 0.04 a | 26.1 ± 0.5 ab |
| *G. violaceus* | 74.6 ± 2.2 a | 138.7 ± 1.9 b | 174.5 ± 4.4 b | 1012.6 ± 68 ab | 11.8 ± 0.1 a | 1.2 ± 0.1 bc | 84.9 ± 0.5 b | 67.6 ± 1.1 b | 1.14 ± 0.01 ab | 1.4 ± 0.2 a | 27.3 ± 0.5 a |
| *O. acuminata* | 60.4 ± 1.9 b | 180.8 ± 3.4 a | 214.3 ± 3.6 a | 1462.2 ± 202.7 a | 9.9 ± 0.1 ab | 1.3 ± 0.2 b | 54.7 ± 0.9 c | 46.2 ± 0.2 c | 0.91 ± 0.02 b | 1.1 ± 0.1 ab | 25.4 ± 0.4 b |
| *Synechococcus* sp. | *48.2 ± 2.6* c | *147.2 ± 2.3* b | *224.0 ± 12.1* a | *631.6 ± 31.3* b | *8.7 ± 0.6* b | *0.8 ± 0.1* c | *57.5 ± 4.5* c | *38.0 ± 3.3* c | *1.19 ± 0.09* ab | 0.8 ± 0.1 b | *22.7 ± 0.5 c* |
| *T. aestivum* (control) | 96.5 ± 0.9 | 12.0 ± 0.2 | 20.1 ± 0.5 | 386.6 ± 31.1 | 3.1 ± 0.1 | 1.1 ± 0.1 | 257.9 ± 4 | 153.3 ± 4.7 | 2.76 ± 0.04 | 17.7 ± 1.3 | - |

Rubisco CO_2_/O_2_ specificity factor (S_c/o_); Michaelis–Menten semi–saturation constant for CO_2_ at 0 % O_2_ (K_c_) and at 21 % O_2_ ($K_{c}^{21 \% O_{2}})$; Michaelis–Menten semi–saturation constant for O_2_ (K_o_); carboxylation turnover rate ($k_{\mathrm{cat}}^{c})$; oxygenation turnover rate ($k_{\mathrm{cat}}^{o})$; ratios between carboxylation and oxygenation turnover rates and Michaelis–Menten constants represent the carboxylation and oxygenation efficiencies, respectively ($k_{\mathrm{cat}}^{c}$ /K_c_, $k_{\mathrm{cat}}^{c}$/$K_{c}^{21\% O_{2}}$, and $k_{\mathrm{cat}}^{o}$/K_o_); percentage of Rubisco relative to total soluble protein (% Rubisco to TSP); and *in vitro* Rubisco isotope fractionation (ε_Rubisco_). Different letters indicate significant differences among strains. One-way ANOVA followed by Tukey’s post hoc test pairwise comparison for parametric data or Kruskal–Wallis test followed by Dunn’s post hoc test pairwise comparison with Bonferroni adjustment for non–parametric data. Z-test with Holm adjustment for ε_Rubisco_ data. Numbers in italics indicate kinetic data from *C. thermalis* and *Synechococcus* sp. PCC6301 taken from Aguiló-Nicolau et al. (2023). *Triticum aestivum* was included as a control for Rubisco kinetics. Values are means of 3-5 replicates.

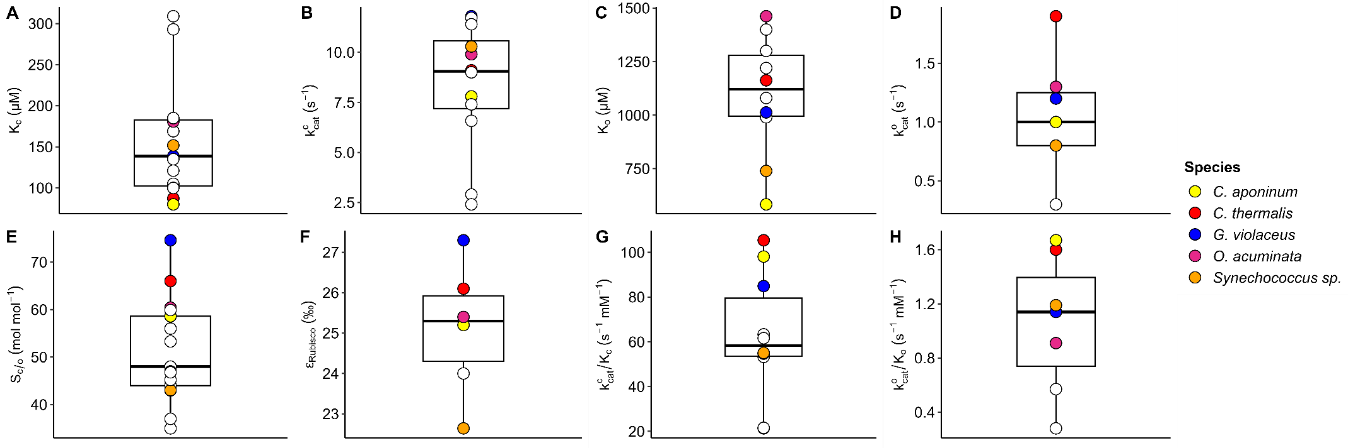

**Figure 1**. Boxplot distribution of Rubisco kinetic parameters in Cyanobacteria, including the analysed strains in the present study (filled circles) as well as previously measured cyanobacterial strains—data compilation from Iñiguez et al. (2020) in empty circles. *C. thermalis* and *Synechococcus* *sp*. PCC6301 data were obtained from Aguiló-Nicolau et al. (2023); except for ε_Rubisco_ which corresponds with original data measured in the present study. Data for other cyanobacterial strains not analysed in the present study are means of the compiled studies for the same strain (Supplementary Spreadsheet 1). **A** K_c_; **B** $k_{\mathrm{cat}}^{c}$; **C** K_o_; **D** $k_{\mathrm{cat}}^{o}$; **E** S_c/o_; **F** ε_Rubisco_; **G** $k_{\mathrm{cat}}^{c}$ /K_c_; **H** $k_{\mathrm{cat}}^{o}$ /K_o_.

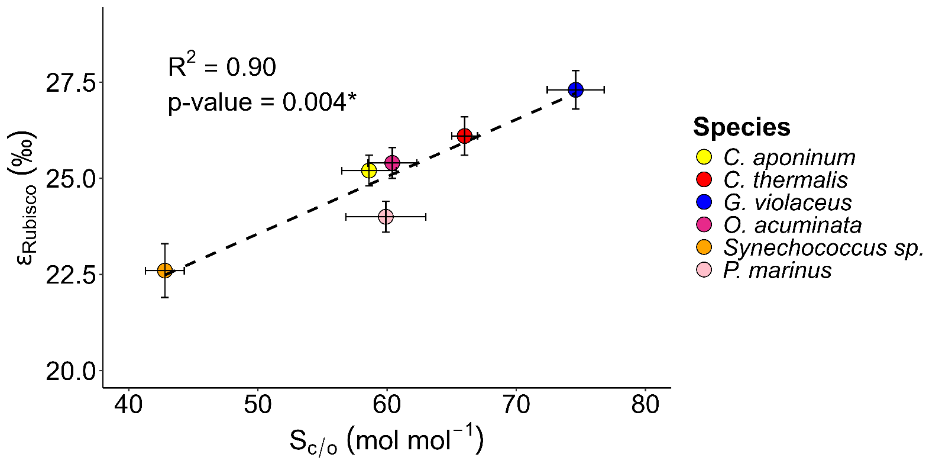

**Figure 2.** Correlation between S_c/o_ and ε_Rubisco_ among Cyanobacteria. *C. thermalis* S_c/o_ value was obtained from Aguiló-Nicolau et al. (2023). *Synechococcus* sp. PCC6301 S_c/o_ and ε_Rubisco_ values are means from this study and other available data (Supplementary Spreadsheet 1). *Prochlorococcus marinus* S_c/o_ value was obtained from Shih et al. (2016) and ε_Rubisco_ from Scott et al. (2007). Values are means ± standard error of 3-5 replicates.

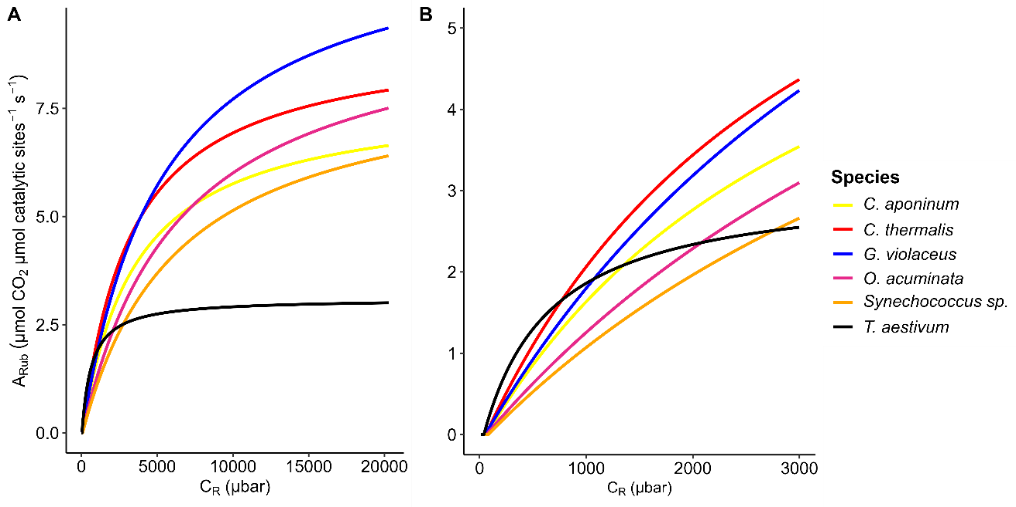

**Figure 3.** Modelled Rubisco-limited assimilation rate (A_Rub_) at 21 % O_2_ and 25 °C at different CO_2_ partial pressures at the Rubisco active site (C_R_), applying the cyanobacterial Rubisco kinetic parameters obtained in the present study. *Synechococcus* sp. and *C. thermalis* kinetic parameters were obtained from Aguiló-Nicolau et al. (2023). *Triticum aestivum* was added as reference using Rubisco kinetic parameters from Table 1. **A** C_R_ ranging from 0 to 20,000 μbar, **B** C_R_ ranging from 0 to 3,000 μbar.

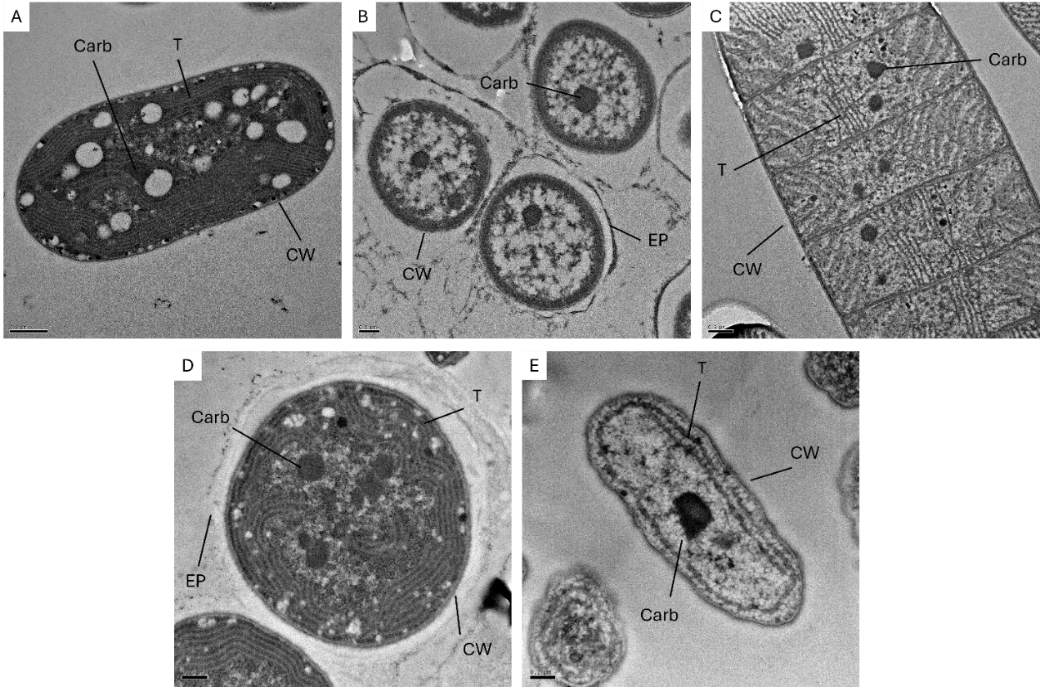

**Figure 4**. Transmission electron microscopy images of **A** *C. aponinum*, **B** *G. violaceus*, **C** *O. acuminata*, **D** *C. thermalis*, **E** *Synechococcus* sp. *Carb* carboxysome, *T* thylakoid membrane, *CW* cell wall, *EP* exopolysaccharide. Scale bars represent: A & C, 500 nm; B, D & E, 200 nm.

**Table 2.** Anatomical differences among cyanobacterial strains (mean ± standard error).

| **Species** | **CellA (μm^2^)** | **NCarb** | **MCarbA (nm^2^)** | **TCarbA (nm^2^)** | **TCarbA/CellA (%)** |
| --- | --- | --- | --- | --- | --- |
| *C. aponinum* | 5.0 ± 0.2 ab | 1.9 ± 0.1 b | 40.0 ± 2.0 c | 74.0 ± 5.0 b | 1.6 ± 0.1c |
| *C. thermalis* | 3.8 ± 0.2 b | 2.9 ± 0.2 a | 31.0 ± 2.0 bc | 82.0 ± 6.0 ab | 2.3 ± 0.2 ab |
| *G. violaceus* | 1.2 ± 0.1 c | 1.1 ± 0.1 c | 32.0 ± 2.0 b | 36.0 ± 2.0 c | 2.7 ± 0.1a |
| *O. acuminata* | 6.3 ± 0.2 a | 1.6 ± 0.1 b | 76.0 ± 3.0 a | 112.0 ± 6.0 a | 1.8 ± 0.1 bc |
| *Synechococcus* sp. | 1.5 ± 0.1 c | 1.8 ± 0.1 b | 20.0 ± 1.0 d | 44.0 ±10.0 c | 2.1 ± 0.1 b |

*CellA* cell area; *NCarb* number of carboxysomes per cell; *MCarbA* average area of carboxysomes; *TCarbA* total carboxysome area per cell; *TCarbA/CellA* percentage of carboxysome area to cell area. Different letters indicate significant differences among strains (One–way ANOVA followed by Tukey’s post hoc test pairwise comparison for parametrical data and Kruskal–Wallis test followed by Dunn’s post hoc test pairwise comparison with Bonferroni adjustment for non–parametrical data). *Chroococcidiopsis thermalis* data were obtained from Aguiló-Nicolau et al. (2023). Results are means of 15-30 replicates.

**Table 3.** Percentage of inhibition of the net photosynthetic rate (A_n_) at 25 °C after the addition of the inhibitors acetazolamide (AZ), ethoxyzolamide (EZ) and 4,4′-diisothiocyanatostilbene-2,2′-disulfonate (DIDS). Mean ± standard error.

| **Species** | **% inhibition AZ** | **% inhibition EZ** | **% inhibition DIDS** |
| --- | --- | --- | --- |
| *C. aponinum* | 4.3 ± 0.3 bc ^*^ | 10.2 ± 0.4 c ^*#^ | 12.4 ± 0.7 b ^*^ |
| *C. thermalis* | 6.3 ± 0.3 ab | 25.2 ± 1.2 abc ^*#^ | 22.0 ± 1.2 a ^*^ |
| *G. violaceus* | 0.2 ± 0.1 c | 30.4 ± 1.1 ab ^*#^ | 12.3 ± 1.3 b ^*^ |
| *O. acuminata* | 7.4 ± 0.5 ab | 21.4 ± 0.6 bc ^*#^ | 6.7 ± 0.8 c ^*^ |
| *Synechococcus* sp. | 11.6 ± 0.6 a ^*^ | 43.4 ± 1.7 a ^*#^ | 14.6 ± 0.7 b ^*^ |

Different letters indicate significative difference among strains (p < 0.05, One-way ANOVA followed by Tukey’s test for parametric data and Kruskal–Wallis followed by Dunn test with Bonferroni adjustment for non–parametric data). In AZ and EZ treatments, an asterisk (*) indicates significant inhibition of the net photosynthetic rate (A_n_), and a hash (#), indicates significant difference in inhibition between the two inhibitors (p < 0.05, repeated measures ANOVA followed by estimated marginal means test with Tukey’s adjustment). In DIDS, asterisk (*) indicates significant inhibition of A_n_ (p<0.05, paired t–test). Results are means of 5-18 replicates.

**Table 4**. *In vivo* photosynthetic semi–saturation constant for the CO_2_ and maximum net photosynthetic rate (A_n max_) at 25 °C obtained from the fitting of the photosynthesis–CO_2_ curves to the Michaelis–Menten equation; and CCM effectiveness, calculated as the ratio between *in vitro* Rubisco $K_{c}^{21\% O_{2}}$ and K_m_ _in vivo_. Mean ± standard error.

| **Species** | **K_m_ _in vivo_ (μM)** | **A_n max_ (μmol O_2_ mg^-1^ DW h^-1^)** | **CCM effectiveness (**$\mathbf{K}_{\mathbf{c}}^{\boldsymbol{21\%}\mathbf{O}_{\mathbf{2}}}\boldsymbol{/}$**K_m_ _in vivo_)** |
| --- | --- | --- | --- |
| *C. aponinum* | 1.3 ± 0.1 c | 1.1 ± 0.2 ab | 92.0 ± 6.6 b |
| *C. thermalis* | 0.5 ± 0.1 e | 0.6 ± 0.1 bc | 212.9 ± 27.5 a |
| *G. violaceus* | 2.7 ± 0.1 a | 0.4 ± 0.1 c | 65.0 ± 4.4 c |
| *O. acuminata* | 0.9 ± 0.1 d | 0.8 ± 0.1 ab | 243.8 ± 18.8 a |
| *Synechococcus* sp. | 2.2 ± 0.1 b | 2.7 ± 0.5 a | 101.9 ± 7.0 b |

Different letters indicate significant differences among species. One–way ANOVA followed by Tukey’s post hoc test pairwise comparison for parametric data and Kruskal–Wallis followed by Dunn test with Bonferroni adjustment for non–parametric data for K_m_ _in vivo_ and A_n max_. Student’s t–test was used for comparing CCM effectiveness means. Results are means of 4-8 replicates.

**SUPPLEMENTARY INFORMATION**

**Supplementary Spreadsheet 1**. Data compilation of Rubisco kinetic parameters from Cyanobacteria (obtained from Iñiguez et al., 2020) plus the data from the present study in “Data” sheet. Information of the species phylogenetic group, type of carboxysomes, principal habitat and the capacity to fix nitrogen in “Strain info” sheet. Bibliographical information in “References” sheet.

**Supplementary Table 1.** Pitman estimator ± standard error and intervals of confidence (CI) at 95 % of ε_Rubisco_ calculated as in Scott et al. (2004).

| **Species** | **Experiment** | **ε_Rubisco_ (‰) ± sd** | **ε_Rubisco_ Pitman estimator (‰) ± se** | **95 % CI for ε_Rubisco_ (‰)** |
| --- | --- | --- | --- | --- |
| *C. aponinum* | 1 | 26.2 ± 0.6 | 25.2 ± 0.2 | 24.5 – 25.9 |
|  | 2 | 25.6 ± 0.4 |  |  |
|  | 3 | 24.3 ± 0.3 |  |  |
| *C. thermalis* | 1 | 27.0 ± 0.9 | 26.1 ± 0.2 | 25.2 – 27.0 |
|  | 2 | 24.6 ± 1.9 |  |  |
|  | 3 | 26.0 ± 0.4 |  |  |
| *G. violaceus* | 1 | 27.3 ± 1.4 | 27.3 ± 0.2 | 26.4 – 28.2 |
|  | 2 | 28.6 ± 2.0 |  |  |
|  | 3 | 27.2 ± 0.4 |  |  |
| *O. acuminata* | 1 | 27.3 ± 0.5 | 25.4 ± 0.2 | 24.8 – 26.29 |
|  | 2 | 24.8 ± 0.6 |  |  |
|  | 3 | 25.8 ± 0.5 |  |  |
| *Synechococcus* sp. | 1 | 23.4 ± 0.7 | 22.7 ± 0.2 | 21.8 – 23.6 |
|  | 2 | 22.6 ± 0.7 |  |  |
|  | 3 | 22.2 ± 0.7 |  |  |

Different values of ε_Rubisco_ for each experiment and each strain are shown together with the standard deviation of the regression applied to calculate each value of ε_Rubisco_. Pitman estimator has been calculated using MATLAB script available in Scott et al. (2004) and assumed as ε_Rubisco_ mean for each strain. 95% confidence intervals obtained when calculating Pitman estimator are displayed.

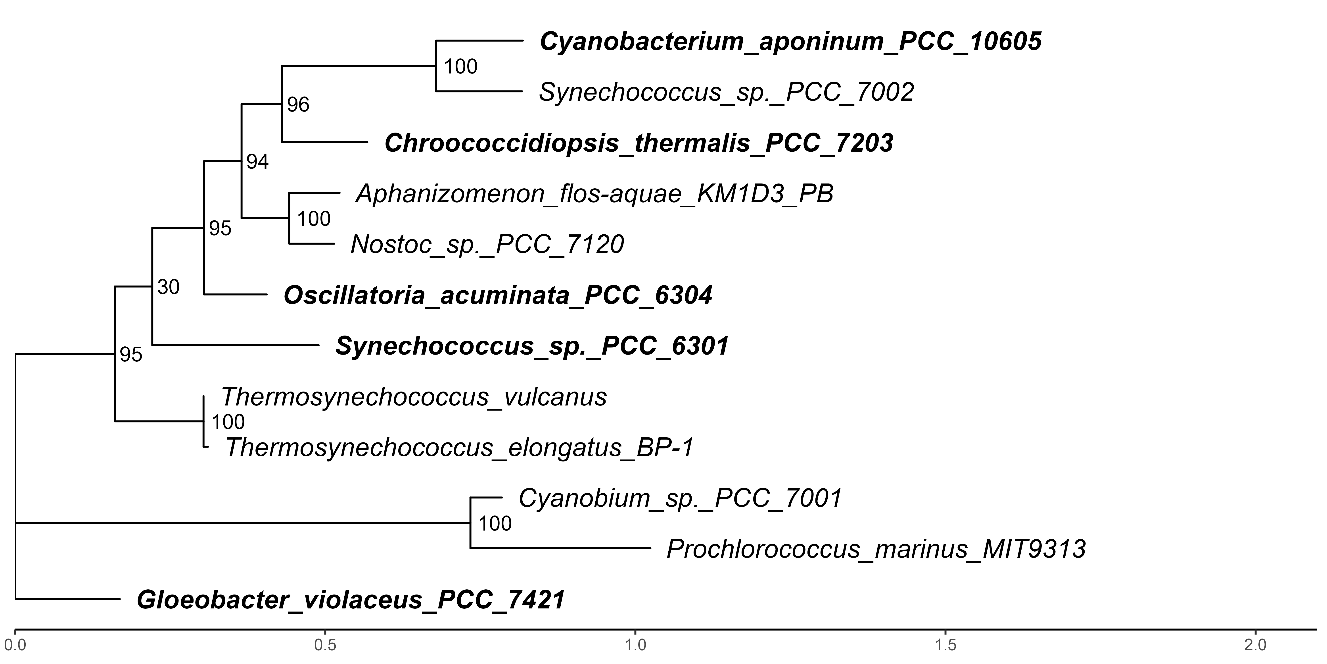

**Supplementary Figure 1**. Maximum likelihood tree inferred from the *rbc*L + *rbc*S of the species in Supplementary Spreadsheet 1. Bootstrap values (from 1000 replicates) are indicated at the nodes. Accession numbers and reference list are provided in Supplementary Table 1. In bold text indicated the species from the present study.

**Supplementary Table 2**. Accession numbers and reference list from the genes used to generate Supplementary Figure 1.

| **Species** | **Accession number** | **Reference** |
| --- | --- | --- |
| *Cyanobacterium aponinum* (PCC 10605) | CP003947.1 | **Gugger M, Coursin T, Rippka R, Tandeau de Marsac N, Huntemann M, Wei C-L, Han J, Detter JC, Han C, Tapia R, Teshima H, Chen A, Krypides N, Mavromatis K, Markowitz V, Szeto E, Ivanova N, Ovchinnikova G, Pagani I, Pati A, Goodwin L, Peters L, Pitluck S, Woyke T, Kerfeld C (n.d.)** Finished genome of chromosome of *Cyanobacterium* sp. PCC 10605. **Unpublished**. |
| *Chroococcidiopsis thermalis* (PCC 7203) | CP003597.1 | **Gugger M, Coursin T, Rippka R, Tandeau de Marsac N, Huntemann M, Wei C-L, Han J, Detter JC, Han C, Tapia R, Davenport K, Daligault H, Erkkila T, Gu W, Munk ACC, Teshima H, Xu Y, Chain P, Chen A, Krypides N, Mavromatis K, Markowitz V, Szeto E, Ivanova N, Mikhailova N, Ovchinnikova G, Pagani I, Pati A, Goodwin L, Peters L, Pitluck S, Woyke T, Kerfeld C (n.d.)** Finished chromosome of genome of *Chroococcidiopsis thermalis* PCC 7203. **Unpublished**. |
| *Gloeobacter violaceus* (PCC7421) | BA000045.2 | **Nakamura Y, Kaneko T, Sato S, Mimuro M, Miyashita H, Tsuchiya T, Sasamoto S, Watanabe A, Kawashima K, Kishida Y, Kiyokawa C, Kohara M, Matsumoto M, Matsuno A, Nakazaki N, Shimpo S, Takeuchi C, Yamada M, Tabata S (2003)** Complete genome structure of *Gloeobacter violaceus* PCC 7421, a cyanobacterium that lacks thylakoids. *DNA Res* **10**: 137–145. |
| *Oscillatoria acuminata* (PCC 6304) | CP003607.1 | **Gugger M, Coursin T, Rippka R, Tandeau de Marsac N, Huntemann M, Wei C-L, Han J, Detter JC, Han C, Tapia R, Davenport K, Daligault H, Erkkila T, Gu W, Munk ACC, Teshima H, Xu Y, Chain P, Chen A, Krypides N, Mavromatis K, Markowitz V, Szeto E, Ivanova N, Mikhailova N, Ovchinnikova G, Pagani I, Pati A, Goodwin L, Peters L, Pitluck S, Woyke T, Kerfeld C (n.d.)** Finished chromosome of genome of *Oscillatoria acuminata* PCC 6304. **Unpublished**. |
| *Synechococcus* sp. (PCC 6301) | AP008231.1 | **Maeda S, Sugita C, Sugita M, Omata T (2006)** Latent nitrate transport activity of a novel sulfate permease-like protein of the cyanobacterium *Synechococcus elongatus*. *J Biol Chem* **281**: 5869–5876. |
| *Cyanobium* sp. (PCC 7001) | DS990556.1 | **Lily E, Ferriera S, Johnson J, Kravitz S, Beeson K, Sutton G, Rogers Y-H, Friedman R, Frazier M, Venter JC (2008)** Direct submission. *J. Craig Venter Institute, Rockville, MD, USA*. Submitted 08-Jul-2008. **Direct submission**. |
| *Anabaena* sp. (PCC 7120) | BA000019.2 | **Kaneko T, Nakamura Y, Wolk CP, Kuritz T, Sasamoto S, Watanabe A, Iriguchi M, Ishikawa A, Kawashima K, Kimura T, Kishida Y, Kohara M, Matsumoto M, Matsuno A, Muraki A, Nakazaki N, Shimpo S, Sugimoto M, Takazawa M, Yamada M, Yasuda M, Tabata S (2001)** Complete genomic sequence of the filamentous nitrogen-fixing cyanobacterium *Anabaena* sp. strain PCC 7120. *DNA Res* **8**: 205–213. |
| *Aphanizomenon flos-aquae* (KM1D3_PB) | CP051528.1 | **Dreher TW, Davis EW II, Mueller RS (2021)** Complete genomes derived by directly sequencing freshwater bloom populations emphasize the significance of the genus-level ADA clade within the Nostocales. *Harmful Algae* **103**: 102005. |
| *Prochlorococcus marinus* (MIT 9313) | BX548175.1 | **Rocap G, Larimer FW, Lamerdin J, Malfatti S, Chain P, Ahlgren NA, Arellano A, Coleman M, Hauser L, Hess WR, Johnson ZI, Land M, Lindell D, Post AF, Regala W, Shah M, Shaw SL, Steglich C, Sullivan MB, Ting CS, Tolonen A, Webb EA, Zinser ER, Chisholm SW (2003)** Genome divergence in two *Prochlorococcus* ecotypes reflects oceanic niche differentiation. *Nature* **424**: 1042–1047. |
| *Synechococcus* sp. (PCC 7002) | CP000957.1 | **Li T, Zhao J, Zhao C, Liu Z, Zhao F, Marquardt J, Nomura CT, Persson S, Detter J-C, Richardson PM, Lanz C, Schuster SC, Wang J, Li S, Huang X, Cai T, Yu Z, Luo J, Zhao J, Bryant DA (2008)** Direct submission. *Department of Biochemistry and Molecular Biology, The Pennsylvania State University, University Park, PA, USA*. Submitted 26-Feb-2008. **Direct submission**. |
| *Thermosynechococcus elongatus* (BP-1) | BA000039.2 | **Nakamura Y, Kaneko T, Sato S, Ikeuchi M, Katoh H, Sasamoto S, Watanabe A, Iriguchi M, Kawashima K, Kimura T, Kishida Y, Kiyokawa C, Kohara M, Matsumoto M, Matsuno A, Nakazaki N, Shimpo S, Sugimoto M, Takeuchi C, Yamada M, Tabata S (2002)** Complete genome structure of the thermophilic cyanobacterium *Thermosynechococcus elongatus* BP-1. *DNA Res* **9**: 123–130. |
| *Thermosynechococcus vulcanus* | AB297499.1  AB297498.1 | **Iwaki T, Shiota K, Al-Taweel K, Kobayashi D, Kobayashi A, Suzuki K, Yui T, Wadano A (2008)** Inhibition of RuBisCO cloned from *Thermosynechococcus vulcanus* and expressed in *Escherichia coli* with compounds predicted by Molecular Operation Environment (MOE). *J Biosci Bioeng* **105**: 26–33. |

**Supplementary Table 3.** Bloomberg’s K and associated p–value for each Rubisco kinetic trait considering the phylogenetic relationships between species according to the phylogenetic tree in Supplementary Figure 1. Data for the Rubisco kinetic traits were obtained from the present study and from previous studies as summarized in Supplementary Spreadsheet 1.

| **Trait** | **Bloomberg’s K** | **p-value K** |
| --- | --- | --- |
| $k_{\mathrm{cat}}^{c}$ | 0.5 | 0.77 |
| K_c_ | 0.5 | 0.19 |
| K_o_ | 0.3 | 0.87 |
| S_c/o_ | 0.4 | 0.27 |
| ε | 0.8 | 0.53 |
| $k_{\mathrm{cat}}^{o}$ | 0.5 | 0.65 |
| $k_{\mathrm{cat}}^{c}$/K_c_ | 0.5 | 0.24 |
| $k_{\mathrm{cat}}^{o}$/K_o_ | 0.7 | 0.72 |

**Supplementary Table 4.** Percentage of A_Rub max_ at C_a_ = 400 ppm.

| **Species** | **C_R_ at C_a_ = 400 ppm (ppm)** | **A_Rub_ at C_a_ = 400 ppm**  **(μmol CO_2_ μmol Rubisco active sites^-1^ s^-1^)** | **A_Rub max_ (μmol CO_2_ μmol Rubisco active sites^-1^ s^-1^)** | **Percentage of A_Rub max_ at C_a_ = 400 ppm (%)** |
| --- | --- | --- | --- | --- |
| *C. aponinum* | 36,800 | 7.1 | 7.8 | 91.3 |
| *C. thermalis* | 85,160 | 8.9 | 9.2 | 96.4 |
| *G. violaceus* | 26,000 | 9.8 | 11.8 | 83.3 |
| *O. acuminata* | 97,520 | 9.3 | 9.9 | 93.8 |
| *Synechococcus* sp. | 40,760 | 7.3 | 8.4 | 86.8 |

CO_2_ concentration at Rubisco active sites (C_R_) under ambient CO_2_ (C_a_ = 400 ppm) was calculated as the product of C_a_ and CCM effectiveness (Table 4) for each strain. Liquid-phase Rubisco kinetic parameters were converted to gas-phase using Henry’s Law (CO_2_ solubility of 0.0328 mol L^-1^ Bar^-1^; Galmés et al., 2016). Rubisco assimilation rate (A_Rub_) at C_a_ = 400 ppm was calculated applying the Farquhar’s biochemical model at each C_R_ (see section Modelling Rubisco Assimilation). A_Rub_ when C_R_ tended to infinite was used to estimate the maximum assimilation rate (A_max_). The percentage of A_Rub max_ at C_a_ = 400 ppm was obtained by dividing A_Rub_ and A_max_, multiplied by 100.

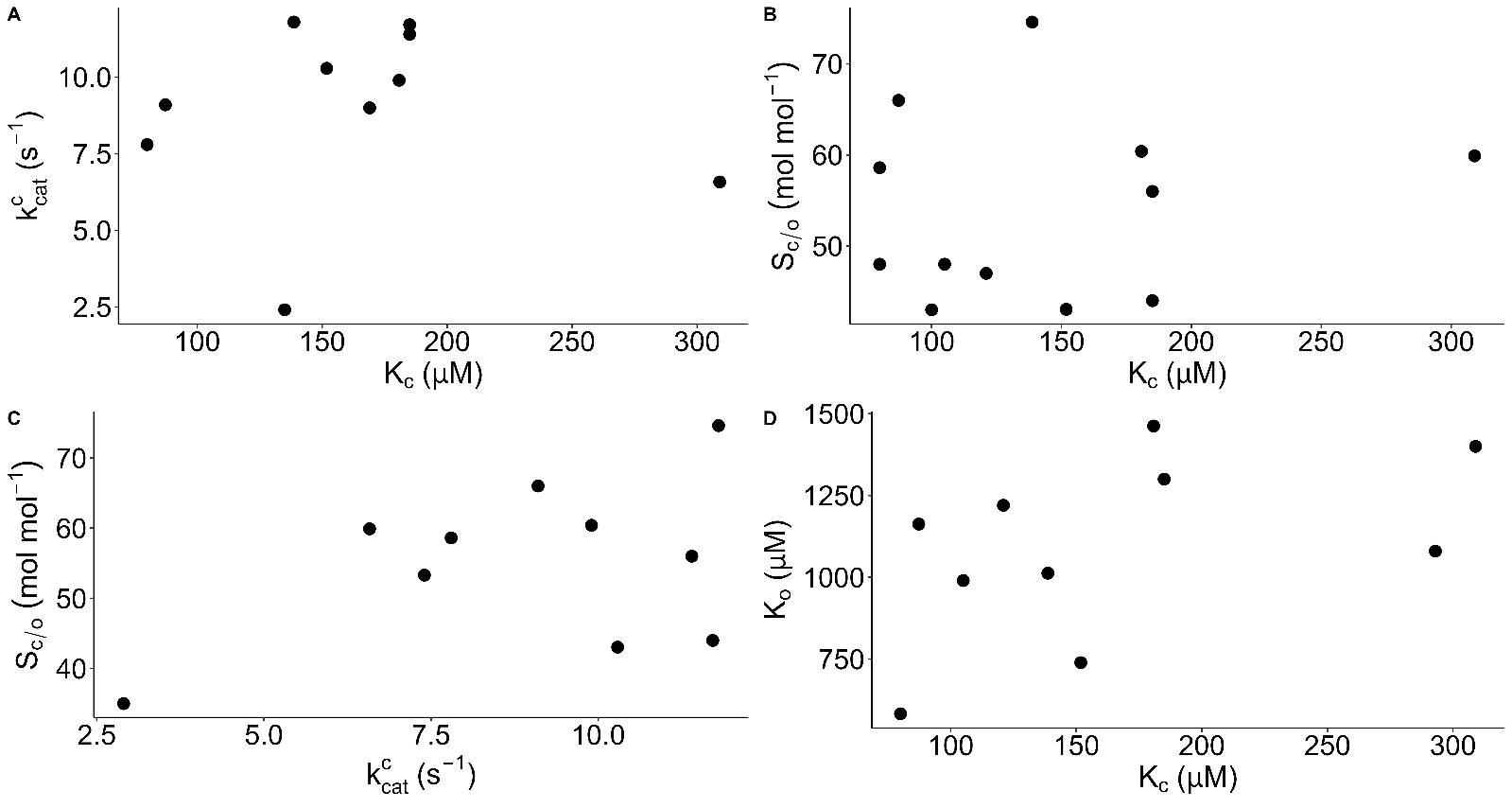

**Supplementary Figure 2**. Correlation plots of **A** $k_{\mathrm{cat}}^{c}$ versus K_c_, **B** S_c/o_ versus K_c_, **C** S_c/o_ versus $k_{\mathrm{cat}}^{c}$, **D** K_c_ versus K_o_. Data was obtained from Supplementary Spreadsheet 1 and the results from the present study.

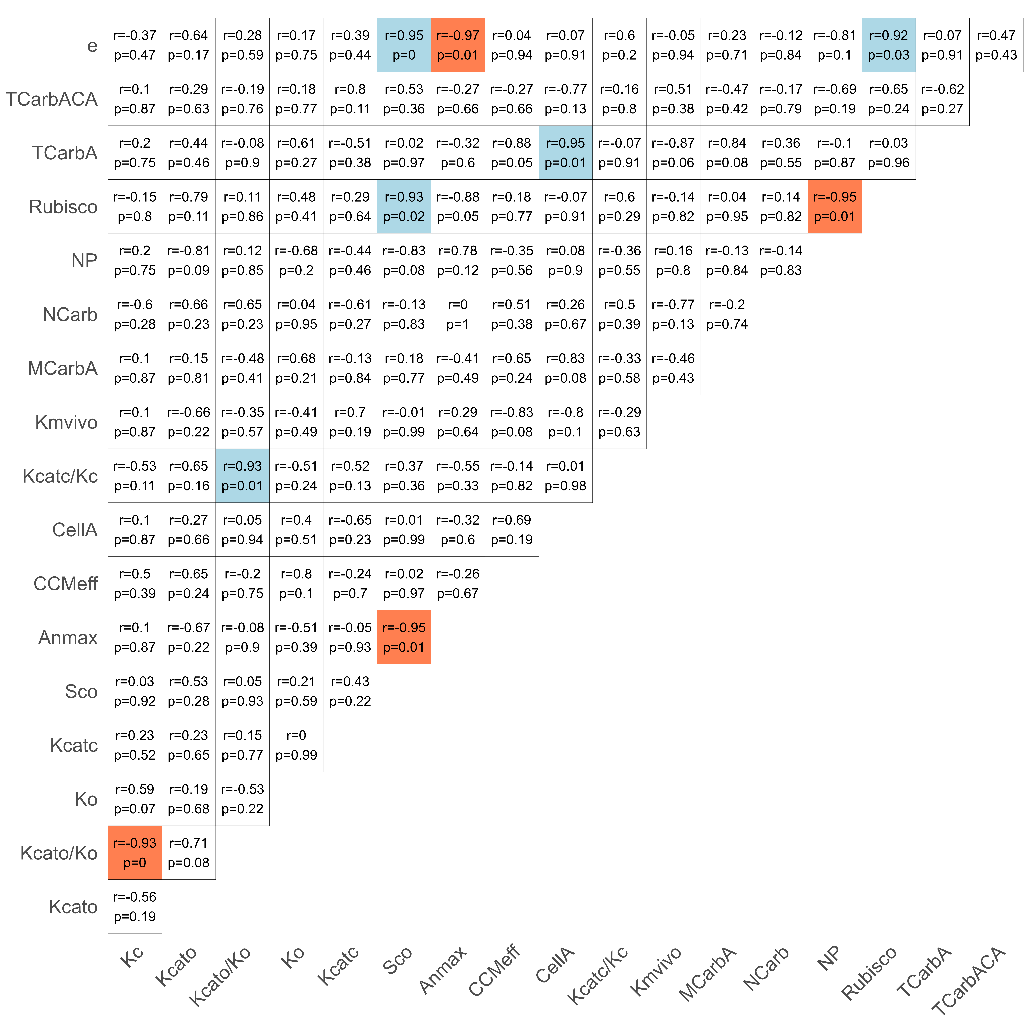

**Supplementary Figure 2**. Correlation matrix of all the measured parameters from cyanobacterial strains, combining data from Iñiguez et al. (2020) and the present study (Supplementary Spreadsheet 1). Coloured squares indicate statistically significant correlations (p–value < 0.05): light blue for positive and light red for negative correlations. Each cell shows the correlation coefficient (r) and the associated p–value (p). Normality was assessed using the Shapiro–Wilk test. Correlations were calculated using the Pitman test for normally distributed variables and Spearman’s test for non–normal distributions. **e**, ε_Rubisco_; **TCarbACA**, total carboxysome area per cell area; **TCarbA**, total carboxysome area; **Rubisco**, Rubisco concentration; **NP**, net photosynthesis; **NCarb**, number of carboxysomes; **MCarbA**, average area of carboxysomes; **Kmvivo**, Michaelis-Menten semi-saturation constant for CO_2_ measured *in vivo*; **Kcatc/Kc**, Rubisco carboxylation efficiency; **CellA**, cell area; **CCMeff**, CCM effectiveness; **Anmax**, maximum net assimilation *in vivo*; **Sco**, Rubisco specificity factor; **Kcatc**, Rubisco carboxylation turnover rate; **Ko**, Rubisco Michaelis-Menten semi-saturation constant for O_2_; **Kcato/Ko**, Rubisco oxygenation efficiency; **Kc**, Rubisco Michaelis-Menten semi-saturation constant for CO_2_; **Kcato**, Rubisco oxygenation turnover rate.
